# Genomic consequences of range expansion and colonisation in the reed warbler (*Acrocephalus scirpaceus*)

**DOI:** 10.1101/2022.11.28.518135

**Authors:** Camilla Lo Cascio Sætre, Katja Rönkä, Mark Ravinet, Severino Vitulano, Matteo Caldarella, Petr Procházka, Bård Stokke, Angélica Cuevas, Nora Bergman, Rose Thorogood, Kjetill S. Jakobsen, Ole K. Tørresen, Fabrice Eroukhmanoff

## Abstract

Range expansion is a common natural phenomenon, which may be intensified by human-induced drivers such as climate change and alterations of habitat. The genetic consequences of range expansion are potentially major, and it is important to study known cases of range expansion to understand how human activities affect contemporary evolution, and to learn more about the genetic adaptive potential of species. The reed warbler (*Acrocephalus scirpaceus*) is a long-distance migratory bird breeding in Eurasia and wintering south of Sahara. It is currently expanding its range northwards, likely as a consequence of climate change. Interestingly, however, reed warblers have also recently colonised new territory southwards, following habitat restoration at the southern range margin. In this study, we investigate the genetic consequences of these two-directional range expansions with RAD-seq, looking at 10 populations from north to south in Europe. We investigate population structure and genome diversity, and assess the role of selection in divergence between populations across the species range. We do not find evidence of strong genetic structure in the reed warbler populations, and the youngest edge populations do not exhibit any substantial loss in genetic diversity, suggesting ongoing gene flow. On a smaller scale, the edge populations are the most genetically distinct, and we identify environmental disparity, especially in precipitation variability, as the main barrier of gene flow, to a greater extent than geographic distance. We find no evidence that the loci involved in population divergence and adaptation in the core populations are the same that are involved in adaptation at the range edges. Using three genome scan methods to identify selection, we found 49 SNPs putatively under selection, of which 33 were located in introns of 28 unique genes. Most of these are correlated with differences in climatic variables of temperature and precipitation. Some genome scan outliers show signs of being part of nascent selective sweeps, especially one which is distinct for the northern range edge. Our results suggest that in the reed warbler, contemporary range expansion has had little effect on molecular diversity and has been rapidly followed by local adaptation to climatic conditions, which could further corroborate the rapid pace at which colonisation of novel environments has occurred both northwards and southwards.

## Introduction

When a species expands its range into new territories, the genetic consequences are potentially major. Founder effects, bottlenecks, and adaptations to the new environment may all shape the genetic composition and population structure of the expanding species (Excoffier et al. 2009; Welles and Dlugosch 2018). Species may expand or shift their range for a number of reasons of which human-induced drivers, such as climate change and habitat destruction/restoration, are of particular interest and different scenarios may lead to very different outcomes. The speed of the expansion, the number of founder individuals, and the amount of gene flow between core and edge populations are among the factors which determine genetic diversity and differentiation in edge populations (Excoffier et al. 2009; Vandepitte et al. 2012; Engler et al. 2016). With small population sizes and high drift, genetic diversity such as heterozygosity, is expected to be reduced at the expansion front, along with an increase in genetic structuring (Austerlitz et al. 1997; Wegmann et al. 2006; Excoffier et al. 2009). Many different empirical studies have demonstrated one or both of these phenomena (Eckert et al. 2008; Vandepitte et al. 2012; Swaegers et al. 2013; White et al. 2013). However, under different circumstances, for instance for highly mobile species capable of long-distance dispersal, genetic diversity may be maintained at range edges, and population structure decreased (Ray and Excoffier 2010). The latter effect was seen in two species of *Hippolais* warblers (Engler et al. 2016), which are long-distance migratory birds with high dispersal propensity. However, there are still only a few case studies that have looked at the genomic consequences of range expansion in species with high dispersal abilities, and the outcome may vary between species. For example, in the highly mobile Indo-Pacific lionfish (*Pterois volitans*), a clear reduction in observed heterozygosity was observed in the expansion front, though population structure was low (Bors et al. 2019).

Local adaptations may be important for the success of the colonisation of new territories, meaning certain loci should diverge between populations. In climate-driven range expansions one may find divergence along climatic gradients and adaptations to the changing climate may occur throughout the range. Genetic signatures of climate-driven selection have been found for instance in damselflies (Dudaniec et al. 2018), marine species (Han and Dong 2020) and birds (Bay et al. 2018; Lamb et al. 2018). In birds, traits like reproductive timing and migration patterns have been shown to be affected by warming temperatures (Gienapp et al. 2007; Koleček et al. 2020). While plasticity may play an important role, such adaptations may also involve genetic changes (Charmantier and Gienapp 2014), as was shown experimentally in the long-distance migratory pied flycatcher (*Ficedula hypoleuca*) (Helm et al. 2019).

Range expansions following habitat restoration may have very different outcomes, depending on the size, distance and environment of the new territory, as well as the historic range and population size of the expanding species. It is crucial to learn more about how human activities affect natural populations, and to what extent different species may be expected to establish and persist in new or altered territories. It is therefore important to study different known cases of range expansions, since the genetic consequences may vary greatly.

The Eurasian reed warbler (*Acrocephalus scirpaceus*) is a long-distance migrant passerine nesting in wetland habitats and wintering in sub-Saharan Africa. The reed warbler is quite unique in that it has simultaneously expanded its range both northwards and southwards in Europe, due to different anthropogenic factors, namely climate change and habitat restoration, respectively. The reed warbler expanded throughout Europe after the Last Glacial Maximum (Arbabi et al. 2014), as temperatures increased and suitable habitat became available, and has successively expanded its distribution towards the north and north-east into Fennoscandia during the 20th century (Järvinen and Ulfstrand 1980; Røed 1994; Stolt 1999; Brommer et al. 2012). Climate change is very likely a major driver for this expansion, as recent changes in temperature have increased productivity in the northern range margin (Eglington et al. 2015; Meller et al. 2018), and there is a strong positive relationship between the presence of reed warblers and temperature in the northern part of Europe (Virkkala et al. 2005; Davies 2019). Reed warblers colonised Finland in the 1920s (Järvinen and Ulfstrand 1980), and had already started expanding in Sweden earlier than that (Stolt 1999). In Norway, the reed warbler was first seen in 1937, and first recorded breeding in 1947 (Røed 1994), and has particularly expanded from the 1970’s (Shimmings and Øien 2015).

The reed warbler has been described as a climate winner (Leisler and Schulze-Hagen 2011). Most reed warbler populations are stable or increasing, unlike many other long-distance migrants (Both et al. 2010; Vickery et al. 2014). Interestingly, in addition to the climate-driven northwards expansion, reed warblers very recently colonised Malta, which is at the southern border of the reed warbler breeding distribution, with a warm and dry climate. There was a lack of suitable habitat there until the early 1990’s, when a wetland area was restored and conserved, containing both Phragmites reed beds and tamarisk groves (Tamarix sp.). From around 1995 and onwards, Malta has sustained a small breeding population of reed warblers. The colonisation of Malta may have been aided by the reed warblers’ adaptive potential. Reed warblers have been shown to respond to warming climate by breeding earlier and longer, thereby increasing productivity (Halupka et al. 2008; Halupka et al. 2021). Local adaptations such as changes in wing shape (Kralj et al. 2010), body mass (Salewski et al. 2010; Sætre et al. 2017) and migratory behaviour (Chamorro et al. 2019) have also been shown to emerge rapidly in reed warbler populations. However, reed warblers may be adversely affected by drought, both at breeding grounds (Jiménez et al. 2018), and along migration routes (Halupka et al. 2017), and some populations are in decline due to reduction of wetland areas and degradation of reed beds (Pollo et al. 2018).

To what extent the range expansions and new climatic selection pressures have influenced the population structure and the genomic composition of the reed warbler is not yet known. Genomic resources for the reed warbler had been lacking until 2021 (Sætre et al. 2021), and most previous studies have focused on ecological questions, at most using a few genetic markers. In studies using microsatellites as genetic markers, genetic differentiation has been found to be generally low between reed warbler populations, but moderate levels of differentiation have been connected to for instance migratory behaviour (Procházka et al. 2011) and wing shape (Kralj et al. 2010). In this study, we examine population structure and genetic patterns among 10 populations in Europe using RAD-sequencing. The selected populations span from the northern to the southern range margins, meaning we are able to compare both directions of range expansion. We determine the role of climatic dissimilarity in genetic divergence, along with geographic distance. Furthermore, we test if local adaptation has left strong signals of selection among the reed warbler populations, by identifying candidate loci under selection, and testing for local adaptation linked to climatic variables. The recently assembled and annotated reed warbler genome (Sætre et al. 2021) enables us to investigate the function of genes associated with candidate regions.

## Materials and Methods

### Sampling

We sampled reed warblers (N = 109) from the following 10 countries from north to south: Finland, Norway, Germany, the Czech Republic (hereafter referred to as Czech), Slovakia, France, Croatia, Italy, Turkey and Malta (Table 1, Figure 1A). We consider Finland and Norway to be the northern edge populations, and Malta the southern edge population, and refer to the remaining seven populations as “core populations” in some analyses. Within Czech, Finland, Norway and Germany we sampled from multiple sites and averaged variables (see Supplementary table 1 for site information). The birds were caught with mist nets, ringed and blood was collected from a brachial vein. The blood samples were stored in either standard Queen’s lysis buffer or ethanol. All birds were released immediately upon completing sampling, and all applicable international, national, and/or institutional guidelines for the care of animals were followed.

**Figure 1.**
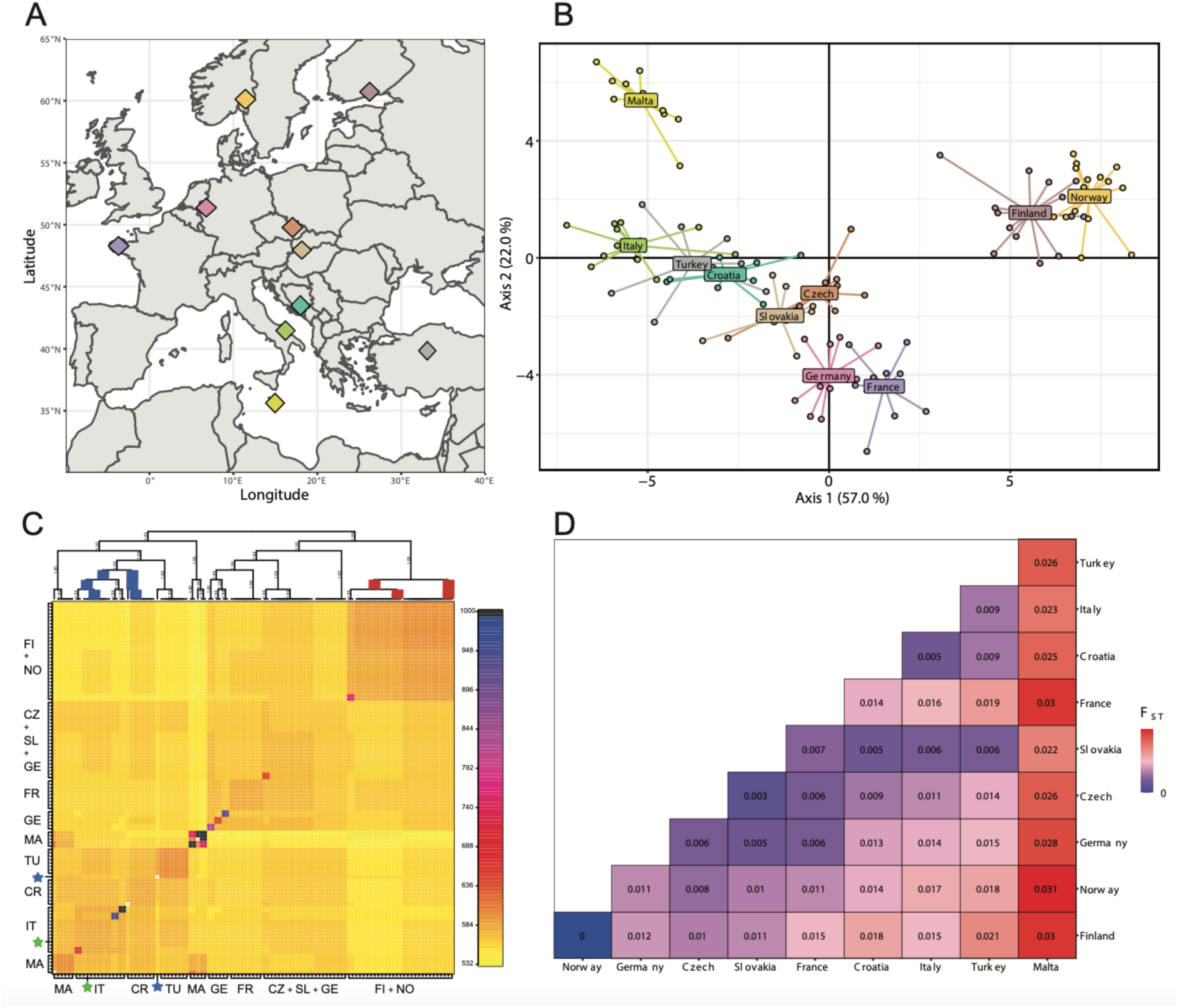
Population structure. A: Geographic position of populations. Map showing the average position of the 10 reed warbler populations as diamonds in different colours, which correspond to the colours in B. B: Population structure based on SNPs. The first two principal axes of a DAPC analysis using site ID as priors. In parentheses, the percentage of variance explained by each axis. C: Population structure based on haplotypes. Population averaged co-ancestry plot with a cladogram from fineRADstructure. The cladogram tree shows the posterior population assignment probabilities of each branch, with 1 having the highest probability (100%). All branches have probabilities of 1, except the branches marked as blue (probability = 0.87) and red (probability = 0.31). The heat map shows coancestry values between pairs of individuals, where the lowest coancestry estimates are towards yellow in colour, and the highest are towards blue and black colour. Each tick on the axes is an individual, and blocks of individuals are indicated by the population names. The stars represent single individuals that have been assigned outside of their population - the green star is an individual from Croatia and the blue star is an individual from Slovakia. D: Population differentiation. Heatmap of pairwise FST between population pairs. The colour gradient goes from blue to red with increasing values of FST, meaning lower FST values are more blue, and higher FST values are more red. Negative values have been set to zero.

**Table 1.** Overview of the populations sampled in this study sorted from highest to lowest latitude, including coordinates, sample size (N) and six selected environmental variables from WorldClim. Longitude and latitude are presented as decimal degrees (rounded to the nearest hundredths). A.Temp = annual mean temperature (℃). Temp.S = temperature seasonality (standard deviation * 100, ℃). M.Temp = maximum temperature of the warmest month (℃). A.Prec = annual precipitation (millimetres). Prec.S = precipitation seasonality (CV, percentage). C.Prec = precipitation of the coldest quarter (millimetres).

| Country | Long. | Lat. | N | A.Temp | Temp.S | M.Temp | A.Prec | Prec.S | C.Prec |
| --- | --- | --- | --- | --- | --- | --- | --- | --- | --- |
| Finland | 60.33 | 25.61 | 14 | 4.62 | 847 | 21.89 | 671 | 28.37 | 152 |
| Norway | 59.69 | 11.45 | 15 | 5.31 | 746 | 20.67 | 789 | 27.28 | 155 |
| Germany | 50.92 | 7.04 | 10 | 9.64 | 588 | 22.74 | 732 | 14.03 | 177 |
| Czech | 49.35 | 16.68 | 10 | 8.76 | 743 | 24.30 | 578 | 44.87 | 84 |
| Slovakia | 48.36 | 17.55 | 10 | 9.83 | 779 | 26.40 | 625 | 31.16 | 111 |
| France | 47.90 | -4.36 | 9 | 12.14 | 361 | 20.60 | 1011 | 36.37 | 311 |
| Croatia | 43.06 | 17.61 | 9 | 16.52 | 722 | 32.00 | 1001 | 31.96 | 302 |
| Italy | 41.54 | 15.89 | 13 | 16.17 | 632 | 30.80 | 442 | 24.43 | 116 |
| Turkey | 39.75 | 32.79 | 8 | 10.93 | 832 | 29.50 | 383 | 41.46 | 123 |
| Malta | 35.95 | 14.38 | 11 | 18.91 | 510 | 30.60 | 516 | 83.23 | 177 |

### DNA extraction, RAD sequencing and bioinformatic processing

DNA was extracted from blood samples using a DNeasy Blood & Tissue Kit (Qiagen, California, USA), following the manufacturers’ protocol. DNA concentration was determined with a Qubit fluorometer and Qubit dsDNA HS assay (Thermo Fisher Scientific, Waltham, MA, USA). Samples were normalised to a concentration of 20 ng/ml, and sent to Floragenex, Inc. (Oregon, USA). DNA was digested with the enzyme *Sbf*I and RAD library construction followed Baird *et al*. (2008). Individual libraries were single-end sequenced on an Illumina HiSeq2000 platform. All samples except the ones from Finland were run on the same plate, generating 742 M reads in total (7.8 M per individual on average). The Finnish samples were run alongside other samples divided over two plates, where plate 1 generated 725 M reads (7.6 M per individual on average) and plate 2 generated 673 M reads (7.1 M per individual on average). Five samples from the first batch were included on the plates with the Finnish samples as controls to check for possible batch effects (see below). Sequence quality was assessed with FastQC (Andrews, 2010). Median Phred score was >30 per base along the length of all reads.

The reads were processed and cleaned using the *process_radtags* pipeline in Stacks v2.5 (Catchen *et al*. 2011, 2013), where low quality reads were discarded (-q filter), reads that contained uncalled bases were removed (-c filter), and barcodes and RAD tags were rescued (-r filter), and the read length were truncated to 91 bp (-t 91). The reads were truncated because the second batch with the Finnish samples produced longer reads than the first batch (141 bp and 91 bp without barcode, respectively), to ensure equal lengths for all samples. The reads were then aligned to the reed warbler reference genome bAcrSci1 (Sætre *et al*. 2021) using bwa mem (Burrows-Wheeler alignment tool, bwa v0.7.17; Li and Durbin 2009) with default options, and sorted with samtools sort (samtools v1.11; Danecek *et al*. 2021). Aligned reads were mapped with the *ref_map* pipeline in Stacks with default options. We ran it first with the five control samples to check for batch effects, and then without the control samples, to use in further analyses.

We produced a SNP dataset with the *populations* pipeline in Stacks, where we only kept loci that were present in at least 80% of individuals within each population (-r 0.80). We also produced a dataset only intended for haplotype analyses, outputted as a RADpainter file (--radpainter), where we filtered haplotype wise (-H flag), including loci present in at least eight populations (-p 8) and at least 80% of individuals across populations (-R 0.80), with a minor allele frequency (--min-maf) of at least 0.05.

The SNP dataset was further filtered with VCFtools v0.1.16 (Danecek *et al*. 2011) and in R v4.0.2 (R core team 2020) step by step (see Supplementary table 2 for SNP count in each step). With VCFtools, we first removed SNPs with a read depth below 10 and above 80, both per genotype (--minDP 10, -- maxDP 80) and per site averaged across individuals (--min-meanDP 10, --max-meanDP 80). We removed SNPs with low read depth as they may be false positive calls, and SNPs with high read depth, as they may be in repetitive regions mapping to multiple parts of the genome. We then removed SNPs found in less than 80% of individuals (--max-missing 0.8), and SNPs with a minor allele frequency under 0.05 (--maf 0.05). In R, we reduced the dataset to one SNP per RAD locus, selecting the SNP with the least amount of missing data, or one at random if the amount of missing data was equal between SNPs. We then split the VCF file into our 10 populations and converted them to BED files with PLINK v1.90b6.13 (Purcell *et al*. 2007). We used these to calculate linkage disequilibrium within each population, outputting pairs of loci with r² values over 0.5 (--r2 --ld-window-r2 0.5), and also to calculate Hardy-Weinberg equilibrium test statistics (--hardy) within each population. In R, we removed SNPs which were in linkage disequilibrium in at least three populations, and SNPs which were in Hardy-Weinberg disequilibrium (p < 0.05) in at least three populations. We also checked A_jk_ relatedness among individual pairs with VCFtools (--relatedness) on the initial SNP dataset from Stacks.

### Controlling for batch effects

To check for possible batch effects between the Finnish samples which were run on two plates in 2021 and the remaining samples which were run on a single plate in 2017, we ran some tests with the five control samples which were included in both years. The control samples were one individual each from the following five populations: Czech, France, Italy, Norway and Turkey. We used the VCF from a *populations* run with all samples and the five controls, without any filters, to run VCFtools with the following flags: --depth, to get the mean depth per individual, --missing-indv, to get missing data per individual, and --het, to get average heterozygosity per individual. We compared the values for the control samples between the 2017 run and the 2021 run, and ran t-tests in R to see if the differences were significant. We also ran bcftools gtcheck (BCFtools v.1.9, Danecek et al. 2021) on the VCF to get pairwise error rates between all individuals, to compare error rates between the control samples from the two batches and the average of the remaining samples. Finally, we ran a principal components analysis (PCA) with PLINK of the control samples to check if they clustered as individuals or as batches.

### Population structure analyses and genetic patterns

From the *populations* run with Stacks, we got estimates of average nucleotide diversity (π), expected and observed heterozygosity and F_IS_ per population in the sumstats_summary output file. We also calculated average allelic richness per population on the VCF from *populations* with the allelic.richness function in hierfstat v0.5-10 (Goudet and Jombart 2021). To see if there were any significant outliers for these variables, we ran Dixon tests with the outliers package v.0.14 (Komsta 2011). For analyses of population structure, we wanted to use unlinked, putatively neutral SNPs, so we used the fully filtered SNP dataset (Supplementary table 2) with 9,236 SNPs. We calculated pairwise F_ST_ values (Weir and Cockerham 1984) among all populations and determined statistical significance with 1,000 permutations using StAMPP v1.6.3 (Pempleton *et al*. 2013) in R.

We further performed various parametric and nonparametric analyses of population structure. We used the adegenet package v2.1.4 (Jombart 2008; Jombart and Ahmed 2011) in R to run the function find.clusters, to compute the goodness of fit (Bayesian Information Criterion, BIC) of each model with an increasing number of clusters (K), and a discriminant analysis of principal components (DAPC). We used cross-validation to find the optimal number of PCs to retain, and site ID as priors. For comparison, we performed a principal components analysis (PCA) with PLINK. We visualised our populations with scatterplots of the first two PCs from both DAPC and PCA with ggplot from the ggplot2 package v3.3.5 (Wickham 2016).

Furthermore, we ran Admixture v1.3.0 (Alexander *et al*. 2009), a model-based analysis which computes maximum-likelihood estimates of parameters to estimate the most likely number of evolutionary clusters. We set the number of assumed populations (K) between 1 and 10, and in order to find the best supported value, we performed a cross-validation analysis. We evaluated the fit of the Admixture models with evalAdmix v0.95 (Garcia-Erill and Albrechtsen 2020).

Finally, using the haplotype dataset, i.e., the RADpainter file outputted by Stacks, we ran fineRADstructure v0.3.2 (Malinsky *et al*. 2018), a model-based Bayesian clustering approach, to infer population structure via shared co-ancestry, with 100,000 burn in iterations (-x), 100,000 sample iterations (-y), sampling every 1,000 (-z). A tree was constructed with 10,000 iterations (-x), and the results were visualised using the scripts fineRADstructurePlot.R and FinestructureLibrary.R available from https://github.com/millanek/fineRADstructure.

### Assessing isolation by distance and environment

Using the fully filtered SNP dataset, we first tested for isolation by distance (IBD) with mantel.rtest from the ade4 package v1.7-17 (Dray and Dufour 2007) with 9999 permutations. We then tested for isolation by distance and isolation by environment (IBE) using multiple matrix regression with randomization (MMRR; Wang, 2013) and variance partitioning via commonality analysis (CA; Prunier *et al*. 2015). The predictor matrices were geographic distance and environmental disparity between all population pairs, and the response matrix was global pairwise F_ST_. We used the distGeo command from the geosphere package v1.5-10 (Hijmans 2019) in R to calculate geographic distances (the geodesic) between the populations, based on the longitude and latitude of each site. To find the environmental disparity between population pairs, we downloaded the 19 bioclimatic predictors (30 s) from WorldClim (Hijmans *et al*. 2005) and used the raster package v3.4-13 (Hijmans 2021) in R to upload them as RasterLayers and extract points from the coordinates of each site, and later take the average values for our 10 populations (see Supplementary materials and Supplementary table 1). To avoid using predictors with high correlation, we only kept the following six for further analyses, which were not highly correlated for our sites (Pearson’s correlation < 0.7, measured in R): Bio 1—annual mean temperature (℃), Bio 4—temperature seasonality (standard deviation * 100, ℃), Bio 5—maximum temperature of the warmest month (℃), Bio 12—annual precipitation (millimetres), Bio 15—precipitation seasonality (CV, percent), and Bio 19—precipitation of the coldest quarter (millimetres). We chose these particular predictors because we consider it relevant to include both measures of temperature and precipitation, and because we wanted to look at both annual trends (annual mean temperature and annual precipitation), measures of variability (temperature and precipitation seasonality), and more extreme measurements (maximum temperature of the warmest month and precipitation of the coldest quarter). We also ran a PCA of these predictors with the prcomp command in R. Data was Z-transformed (i.e., standardised by subtracting the mean and dividing by the standard deviation) to make the regression coefficients of the predictors comparable (Prunier *et al*. 2015).

### Range expansion statistics

To explore the direction of range expansion between population pairs, we used the rangeExpansion package v. 0.0.0.9000 in R (Peter and Slatkin 2013), using the fully filtered SNP dataset. We calculated the directionality index, ψ, for all population pairs with the get.all.psi function. The index ψ detects the allele frequency clines created by successive founder events, and assumes the populations furthest away from the origin of the expansion to have experienced more genetic drift (Peter and Slatkin 2013). Between two populations *i* and *j*, ψ*_ij_* should therefore be negative if *i* is closer to the origin of the expansion than *j*, around zero if they are equally close or no range expansion has occurred, and positive if *i* is further away from the point of origin than *j*. To determine significance, we calculated the standard deviation of the upper triangle of the pairwise ψ matrix excluding the diagonal, and calculated the Z-score for each population pair.

### Studying divergence at range edges compared to range core

To investigate whether divergence within the range core correlates with divergence between the range edge populations, we calculated F_ST_ per locus with the basic.stats function in hierfstat for the core populations (Croatia, Czech, France, Germany, Italy, Slovakia and Turkey) as well as between Malta and Finland, and Malta and Norway, and ran two separate linear regression analyses. For this analysis, we used a semi-filtered dataset with 22,862 SNPs (step 4 in Supplementary table 2) since the fully filtered dataset excludes loci in Hardy-Weinberg and linkage disequilibrium, to include loci which may have been subjected to selection. Average pairwise F_ST_ between core populations was used as the predictor, and pairwise F_ST_ between Malta and Finland, or Malta and Norway was used as the response variable. We also ran a linear regression to check the correlation between pairwise F_ST_ between Malta and Finland and pairwise F_ST_ between Malta and Norway.

### Identifying signatures of selection

To identify candidate loci under selection we ran three different genome scan analyses using a semi-filtered dataset with 22,862 SNPs (step 4 in Supplementary table 2). We ran BayeScan v2.1 (Foll and Gaggiotti, 2008), a Bayesian method based on multinomial-Dirichlet likelihood, that detects loci with high differentiation (F_ST_) compared to the rest of the genome. We ran it with default parameters for the chain (-n 5000 -thin 10 -nbp 20 -pilot 5000 -burn 50000), but increased the prior odds for the neutral model from 10 to 100 (-pr_odds 100) to reduce the number of false positives given the number of SNPs tested. We used PGDSpider v2.1.1.5 (Lischer and Excoffier 2012) to convert the VCF file to a BayeScan input file. We evaluated the convergence of the chain with coda v0.19-4 (Plummer *et al*. 2006). We also ran OutFLANK v0.2 (Whitlock and Lotterhos 2015), which identifies outliers by inferring a distribution of neutral F_ST_ using likelihood on a trimmed distribution of F_ST_ values. We ran MakeDiploidFSTMat to create an OutFLANK input file, using the genotypes from the VCF recoded with vcftools --012. We then ran the OutFLANK function with default parameters (LeftTrimFraction = 0.05, RightTrimFraction = 0.05, Hmin = 0.1, qthreshold = 0.05), evaluating the model by following the procedure explained here: https://rpubs.com/lotterhos/outflank.

Finally, to identify candidate loci under selection that are related to specific environmental variables, we ran BayeScEnv v1.1 (de Villemereuil and Gaggiotti, 2015), which is an extension of BayeScan incorporating environmental data. We used the same environmental variables as used in the MMRR analysis (see above). We ran BayeScEnv separately for each environmental variable with default parameters (-n 5000 -thin 10 -nbp 20 -pilot 5000 -burn 50000 -pr_jump 0.1 -pr_pref 0.5), and set the false discovery rate to 0.05. We evaluated the convergence of the chains with coda v0.19-4. We used the ggVennDiagram package v1.2.1 in R to create a Venn diagram of the outliers detected by the three genome scan methods used. We used the genome annotation of the reed warbler (available on Figshare: https://doi.org/10.6084/m9.figshare.16622302.v1) to find the genomic location of our outlier SNPs, and to investigate the function of genes in close proximity (within 20 kb). To investigate population structuring when only looking at the outlier loci found by the three genome scan methods, we ran a PCA with PLINK, and plotted the first two principal components with ggplot.

We conducted analyses of extended haplotype homozygosity to detect recent selective sweeps in the genome. These analyses require a phased VCF, so we used ShapeIt4 (v4.2.2, Delaneau et al. 2019) to create two different phased VCFs, one using just the population data to phase SNPs and one which was informed by phasing of the heterozygous SNPs of the reference assembly. To phase the reference individual, we mapped the reads that had been used for the reference assembly (Sætre et al. 2021), that is, 10X Genomics linked reads, PacBio Continuous Long Reads and Arima Genomics Hi-C reads to the reference assembly. Long Ranger (v2.2.2, Marks et al. 2019) was used to map the 10X reads; bwa mem (v0.7.17, Li and Durbin 2009) with options -5SPM and samtools (v1.9, Danecek et al. 2021) markdup was used to map the Hi-C reads, and pbmm2 v1.2.1 (https://github.com/PacificBiosciences/pbmm2), which uses minimap2 internally (v2.17, Li 2018) was used to map the PacBio reads. FreeBayes v1.3.1 (Garrison and Marth 2012) was used to call SNPs based on the alignment of the 10X reads. Then we used an approach similar to DipAsm (Garg et al. 2021) to phase SNPs. First HapCUT2 (v1.3.2, Edge et al. 2017) was used with the Hi-C and 10X reads to get a phasing at the chromosome level. This was used as input to WhatsHap (v1.0, Martin et al. 2016) together with PacBio reads to get a fine-grained, chromosome-level phasing. We tested both phased VCFs and compared the results. Below, we display the results of the one based just on population data, but the results did not change qualitatively with the reference based VCF. We split the phased VCF into our 31 chromosomes with bcftools view, and used the R package rehh v.3.2.2

(Gautier and Vitalis 2012; Gautier et al. 2017) to first calculate integrated haplotype homozygosity score (iHS) along each chromosome. We read in the VCFs with the data2haplohh function, setting polarize_vcf to FALSE, as we do not have information about ancestral or derived alleles. We then performed the haplotype genome scan with the scan_hh function, again setting polarized as FALSE, and calculated iHS with the ihh2ihs function with freqbin = 1. The iHS statistic is based on a measure of the extent of haplotype homozygosity surrounding a SNP. Higher iHS scoring SNPs are associated with longer haplotypes and lower neighbouring diversity compared to other SNPs. It is intended to be used within-population, but we combined all populations to find the strongest signals of selection across our populations.

We then conducted an analysis of cross-population extended haplotype homozygosity (XP-EHH), which assesses haplotype differences between two populations at the same SNP and allows detection of recent selection events, in which haplotypes have almost or fully proceeded to fixation. We ran the ies2xpehh function on each population pair for each chromosome from the haplotype genome scan data. We only used the 29 main autosomes, and chromosomes Z and W, not smaller scaffolds.

To assess the correlation between genome scan methods and extended haplotype homozygosity methods, we investigated the iHS and XP-EHH -log_10_ p-values of the outliers found by BayeScan, Outflank and BayeScEnv. One of the 49 outliers found by genome scan methods were located on a smaller scaffold, and was therefore excluded from analyses. Another five outliers had missing values for iHS. Therefore, we selected a random set of 48 loci for XP-EHH and 43 loci for iHS and calculated their average -log_10_ p-value 10,000 times each, plotting the result as histograms, to see where the -log_10_ p-values of our outliers fell on the curves.

Finally, we used the Hi-C data used for the reed warbler genome assembly (Sætre et al. 2021) to look at the fraction of significant interactions of the 49 outlier loci found by BayeScan, Outflank and BayeScEnv. We used the Juicer pipeline (v1.6, Durand et al. 2016) to map and process the Hi-C reads. Hi-C_contact_caller (Ray et al. 2019) was used to determine the significance of interactions between the SNPs and transcription start sites of genes. In addition, we calculated the fraction of significant interactions of 49 random loci 200 times, and plotted the result as a histogram, to compare it with the fraction of significant interactions of our outlier loci.

## Results

### Reference mapping and data filtering

By mapping the RAD sequence reads to the reed warbler reference genome, we obtained 120,500 RAD loci with 704,909 SNPs. The mean effective per-sample coverage was 50.9x (min = 23.1x, max = 112.4x). The SNP dataset was filtered further step by step (see Supplementary table 2 for SNP count in each step), with the most stringently filtered dataset containing 9,236 SNPs. For the haplotype dataset, we retained 60,952 RAD loci.

### Controlling for batch effects

We used five control samples which were run both in 2017 with all samples except the samples from Finland, and in 2021 with the Finnish samples to check for possible batch effects. The samples had lower mean depth and a lower frequency of missing data in 2021 than in 2017 (Supplementary table 3), but the differences were not significant with a t-test (t = 1.71, p = 0.14 and t = 2.56, p = 0.06, respectively). However, the samples had higher mean heterozygosity in 2021 than in 2017, a difference which was significant with a t-test (t = -3.58, p = 0.02), though not highly significant. The average (± standard deviation) in 2017 was 0.068 (± 0.009) and in 2021 it was 0.084 (± 0.003). In the PCA of only the control samples from both batches, the samples clustered as individuals, not by batch (Supplementary figure 1).

### Population structure and genetic patterns

Finland has the highest observed heterozygosity and the lowest F_IS_ of all 10 populations (Supplementary table 4). However, this is likely due to some small batch effect, as the control samples had a significantly higher proportion of heterozygotes in the 2021 plates than they had in the 2017 plate, and Finland had a lower average than four of the five control samples in the 2021 plates (higher than the Norwegian control sample). Slovakia has the highest nucleotide diversity and allelic richness, and the highest expected heterozygosity, but fewer observed heterozygosity, and thus the highest F_IS_ (Supplementary table 4). Norway has the second highest F_IS_, the lowest observed heterozygosity, and second lowest nucleotide diversity, and France has the lowest nucleotide diversity and the lowest expected heterozygosity (Supplementary table 4). However, running a Dixon test for detecting an outlier in a sample (either the highest or the lowest value), there are no significant outliers for any of the statistics.

We find consistent support for weak population structure among our 10 populations. The clustering procedure used in DAPC supported the existence of one single cluster (K = 1) within the data. BIC was 986.98 for K = 1, and increased steadily with an increasing number of K (BIC = 1041.19 for K = 40). With Admixture, K = 1 was again the optimum value of K identified by cross-validation error. Using site ID as priors, the scatterplot of the first two principal axes from the DAPC analysis indicates a gradual separation of northern and southern populations following Axis 1 leftwards, with our most northern populations (Finland and Norway) and our most southern (Croatia, Italy, Turkey and Malta) on either end of the axis (Figure 1B). Malta separates from the others on Axis 2, on the opposite end from Germany and France. Discriminant analysis maximises the separation between groups while minimising variation within-group, and we see in the initial PCA plot that a trio and a duo of individuals from Malta clustered far away from the other samples (Supplementary figure 2A). These samples had high A_jk_ relatedness scores with each other within each cluster (A_jk_ > 0.3), so to explore the remaining PCA structure, we removed two individuals from the trio and one from the duo. This PCA scatterplot looks very similar to the DAPC scatterplot, except Malta is not separate from the southern populations (Supplementary figure 2B).

**Figure 2.**
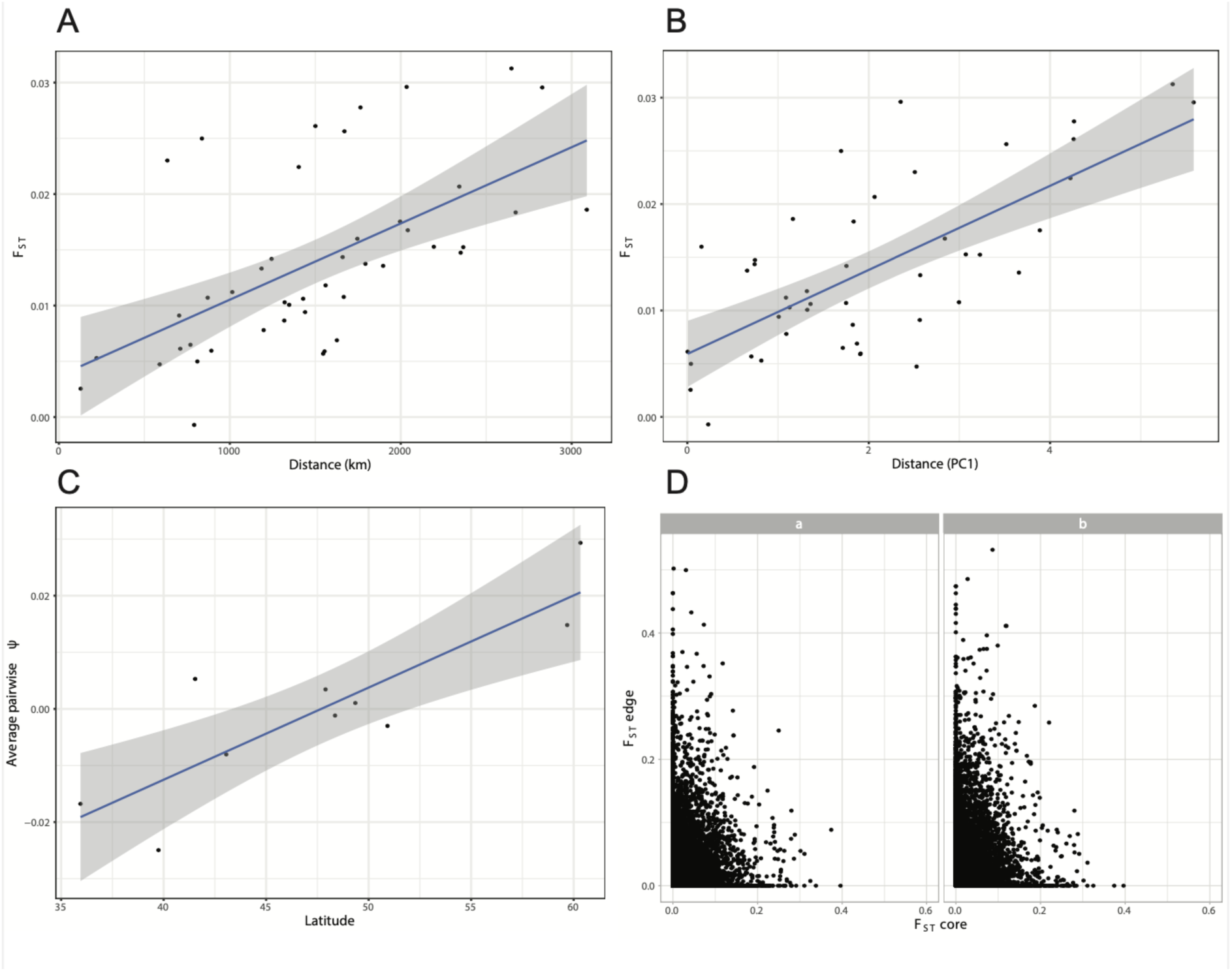
Genome-wide consequences of range expansion. A: Isolation by distance. Pairwise FST against geographic distance in km between each population pair (linear regression: R^2^ = 0.35, p = 2e-05). B: Isolation by environment. Pairwise FST against environmental disparity (absolute difference in PC1 of a PCA of all six environmental parameters) between each population pair (linear regression: R^2^ = 0.48, p = 1e-07). C: Source-sink dynamics in range expansion. Average values per population of pairwise Ψ (directionality index) against latitude (linear regression: R^2^ = 0.73, p = 0.002). D: Novel vs. standing genetic variation between range edges and range core. Average pairwise FST values between the core populations, i.e., Croatia, Czech, France, Germany, Italy, Slovakia and Turkey against pairwise FST between Malta and Finland (a) and Malta and Norway (b). Negative FST values have been set to zero.

Plotting the Admixture results for K = 2 with the populations arranged from higher to lower latitudes (Supplementary Figure 3A), we observe an almost gradual transition in the ancestral fractions from our northernmost to our southernmost populations. However, evaluation of the admixture model with evalAdmix showed a positive correlation of residuals between some individuals within Germany, Italy, Malta and Norway, and both positive and negative correlation between some populations (Supplementary figure 3B), suggesting the model fit is not optimal.

**Figure 3.**
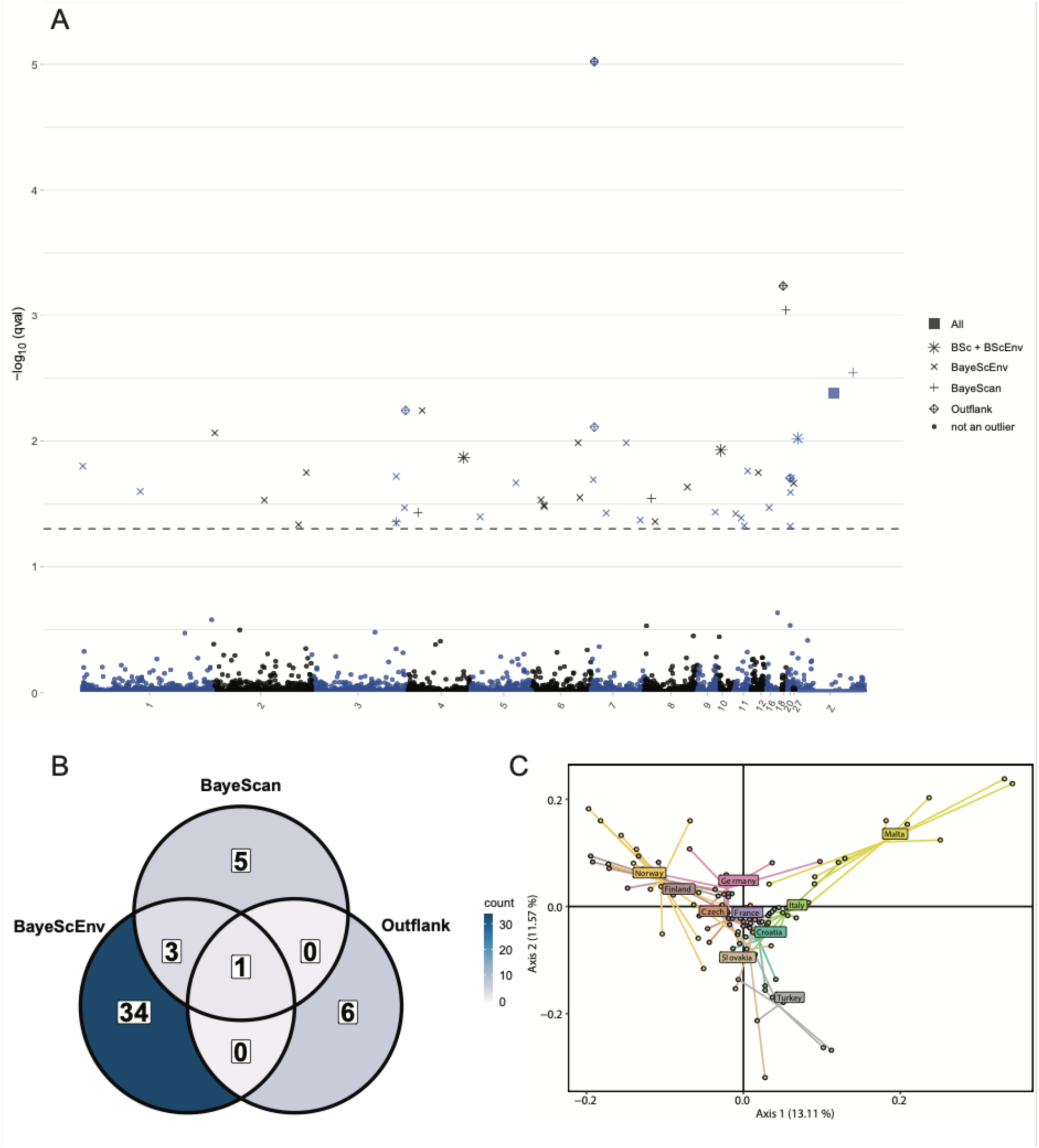
Signatures of selection. A: Outliers across the genome. Manhattan plot of the outliers found by BayeScan, BayeScEnv and Outflank. Only chromosomes with at least one outlier have been included for illustration purposes. The colour of the points shows which chromosome they are in (alternating blue and black), and the shape of the points reflect their outlier status: circles are not outliers, crosses are outliers found only by BayeScEnv, plus signs are outliers found only by BayeScan, diamond plus symbols are outliers found only by Outflank, star symbol found by both BayeScan and BayeScEnv, and square symbol is the outlier found by all three methods. The stapled horizontal line is the threshold for being marked as an outlier (q-value = 0.05). B: Count of outliers found by each method. Venn Diagram of the outliers found by BayeScan, Outflank and BayeScEnv. C: Population structure of outlier loci. The first two principal axes of a PCA analysis, including only the 49 outlier SNPs found by BayeScan, BayeScEnv and Outflank. The percentage of variance explained by each axis are in parentheses.

The population averaged co-ancestry plot with a cladogram from fineRADstructure (Figure 1C) again largely separates the northern populations (Finland, Norway, Germany, Czech, Slovakia and France) and the southern populations (Croatia, Italy, Turkey and Malta). Within the northern group, Finland and Norway separate from the others. Individuals with high levels of co-ancestry within a population tend to form separate groups, which is especially apparent with Malta. Most of the branches in the cladogram tree have very high support (>= 99%), but some branches separating individuals from Croatia and Italy have slightly lower support (87%) and some branches separating individuals from Finland and Norway have quite low support (31%; Figure 1C).

Pairwise F_ST_ ranged from -0.0007 (between Finland and Norway) to 0.031 (between Malta and Norway). The highest F_ST_ values generally included Malta (Figure 1D). The mean (± standard deviation) pairwise F_ST_ was 0.014 ± 0.008. All p-values were highly significant (p < 0.005), except between Finland and Norway (p = 0.79), and Czech and Slovakia (p = 0.03), but the F_ST_ values were still too low (< 0.10) for reliable estimation of migration rates (Meirmans, 2014).

### Isolation by distance and environment

There is significant isolation by distance in our populations according to a Mantel test (p = 0.0002), and a scatterplot of F_ST_ against geographic distance shows that F_ST_ values increase with geographic distance (Figure 2A, linear regression: R_2_ = 0.35, p = 2e-05). However, in a multiple matrix regression analysis, with geographic distance and all six selected bioclimatic predictors, geographic distance is not significant (β = 0.36, p = 0.09), and the commonality analysis shows the unique effect is small (U = 0.04), compared to the common effect (C = 0.31, Table 5). Geographic distance alone thus explains 4% of the total variance in F_ST_. Of the 86% of variance in F_ST_ that is explained by the model (R^2^), geographic distance accounts for around 40% of that variance in total, but most of this is due to common effects, and alone it only accounts for about 5% (Table 5). The most important predictor in this model is Prec.S (precipitation seasonality), which alone explains 34% of the variance in F_ST_, and almost 40% of the variance in R^2^ (Table 5). We see that Prec.S and pairwise F_ST_ are clearly positively correlated in a scatterplot (Supplementary Figure 4, linear regression: R^2^ = 0.53, p = 2e-08). The other bioclimatic variables show very low unique contributions, except A.Temp (annual mean temperature), which explains 2% of the total variance in F_ST_, and around 2% of the variance in R^2^ (Table 5). In another model, with geographic distance and the first principal component of a PCA of all six bioclimatic predictors, PC1 is the most important predictor (β = 0.54, p = 0.02), explaining alone 23% of the total variance in F_ST_, whereas geographic distance (β = 0.35, p = 0.06) explains 10% of the total variance in F_ST_ (Table 6). F_ST_ is clearly higher when the absolute difference in PC1 increases (Figure 2B, linear regression: R^2^ = 0.48, p = 1e-07). These results suggest that divergence between populations is mainly climate-driven.

**Table 5.** Multiple matrix regression with randomization (MMRR) with the Unique (U), Common (C) and Total (T) effects from a Commonality Analysis. Predictor variables are: Geo = geographic distance, A.Temp = annual mean temperature, Temp.S = temperature seasonality, M.Temp = maximum temperature of the warmest month, A.Prec = annual precipitation, Prec.S = precipitation seasonality, C.Prec = precipitation of the coldest quarter. The dependent variable is genetic distance (FST). Beta weights (β) quantify the change in FST (in standard deviation units) with one standard deviation change in the predictor. The Unique effect (U) is the amount of variance in FST that is uniquely accounted for by a predictor. The Common effect (C) is the sum of the variance in FST that can be jointly explained by a predictor and other predictors. The Total effect (T) is the sum of U and C. % U, % C and % T is obtained by dividing U, C and T on the model fit index, R² (respectively) and multiplying with 100. They represent the amount of explained variance in FST (R²) that is accounted for by a predictor, by its unique effects (% U), by its effects shared with other predictors (% C), and by its total effect (% T). Model: FST ∼ Geo + A.Temp + Temp.S + M.Temp + A.Prec + Prec.S + C.Prec. R² = 0.86. p = 0.001.

| Predictor | $\beta$ | p-value | U | C | T | % U | % C | % T |
| --- | --- | --- | --- | --- | --- | --- | --- | --- |
| Geo | 0.36 | 0.09 | 0.04 | 0.31 | 0.35 | 4.61% | 35.75% | 40.36% |
| A.Temp | 0.20 | 0.09 | 0.02 | 0.33 | 0.36 | 2.42% | 39.03% | 41.45% |
| Temp.S | -0.01 | 0.96 | 0.00 | 0.05 | 0.05 | 0.00% | 5.57% | 5.57% |
| M.Temp | 0.11 | 0.39 | 0.01 | 0.16 | 0.17 | 0.63% | 18.94% | 19.57% |
| A.Prec | 0.08 | 0.57 | 0.002 | 0.003 | 0.01 | 0.23% | 0.37% | 0.61% |
| Prec.S | 0.65 | 0.001 | 0.34 | 0.19 | 0.53 | 39.76% | 21.77% | 61.54% |
| C.Prec | -0.02 | 0.88 | 0.0001 | 0.02 | 0.02 | 0.02% | 2.57% | 2.58% |

**Table 6.**
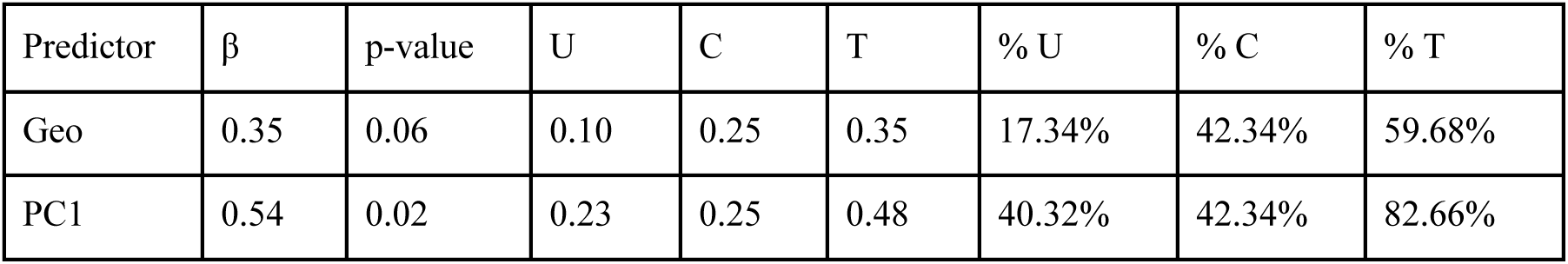
Multiple matrix regression with randomization (MMRR) with the Unique (U), Common (C) and Total (T) effects from a Commonality Analysis. Predictor variables are: Geo = geographic distance, PC1 = first principal component of the PCA of all six bioclimatic variables. The dependent variable is genetic distance (FST). Beta weights (β) quantify the change in FST (in standard deviation units) with one standard deviation change in the predictor. The Unique effect (U) is the amount of variance in FST that is uniquely accounted for by a predictor. The Common effect (C) is the sum of the variance in FST that can be jointly explained by a predictor and other predictors. The Total effect (T) is the sum of U and C. % U, % C and % T is obtained by dividing U, C and T on the model fit index, R² (respectively) and multiplying with 100. They represent the amount of explained variance in FST (R²) that is accounted for by a predictor, by its unique effects (% U), by its effects shared with other predictors (% C), and by its total effect (% T). Model: FST ∼ Geo + PC1. R² = 0.58. p = 0.002.

### Range expansion statistics

To investigate whether the directions of range expansion can be shown with our data, we ran rangeExpansion, which estimates the directionality index, ψ, for all population pairs. Pairwise ψ ranged from -0.03 between Croatia and Finland (i.e., Finland is the sink), and 0.05 between Finland and Turkey (i.e., Finland is the sink). Average values per population of pairwise ψ indicates that the range expansion has gone from south to north, with the southernmost populations on average being the source in pairwise comparisons, and the northernmost being the sink (Supplementary table 5). The central populations have average values very close to zero, meaning that for most of the pairwise comparisons they are almost equally close to the point of origin. There is a significantly positive correlation between average pairwise ψ and latitude (Figure 2C, linear regression: R^2^ = 0.73, p = 0.002). Looking at all 45 pairwise comparisons, five are significant at 5% alpha level according to their Z-scores, all of which include either Norway or Finland.

### Divergence at range edges compared to range core

We compared average F_ST_ values between the seven core populations to F_ST_ values between Malta and Finland and Malta and Norway (Figure 2D), and found that the F_ST_ values are generally higher between both Malta and Finland, and Malta and Norway, than between the core populations, especially at the sites with lower F_ST_ between the core populations. In a linear regression, there is no correlation for Malta–Finland (R^2^ = 0.0001, p = 0.13), and a barely significant correlation for Malta– Norway at a 5% alpha level (R^2^ = 0.0002, p = 0.03), but the extremely low R^2^ values show that divergence between the core populations explain essentially nothing of the variation in divergence between edge populations. This suggests it is not the same loci involved in divergence between edge populations as between core populations. We find that there is a high correspondence between F_ST_ values between Malta and Finland and F_ST_ values between Malta and Norway (linear regression: R^2^ = 0.24, p = <2e-16), suggesting that at least some of the loci involved in divergence between Malta and Fennoscandia are shared between Finland and Norway.

### Identifying signatures of selection

We found 9 outliers with BayeScan, 7 outliers with Outflank, and 38 outliers in total with BayeScEnv scattered across the genome (Figure 3A). In total, we found 49 unique outliers. One of these was found by all three methods, and three were found by two methods (Figure 3B; Supplementary table 7). Thirty-three of the outliers were located in introns of 28 unique genes: 24 protein-coding genes and 4 long non-coding RNAs. The remaining 16 outliers were located in intergenic regions, with seven outliers being close (within 15 kbp) to the nearest gene, two being within 30 kbp to the nearest gene, and seven being over 30 kbp from the nearest gene. Three of the intergenic outliers were close to several genes.

The outlier that was found by all three methods was located in an intergenic region close to (< 15 kbp) the gene *C9orf72*. The protein encoded by this gene is part of a GTPase-interacting complex which is a potent regulator of endocytic transport, autophagy and inflammation, and mutations on this gene is associated with frontotemporal dementia and ALS in humans (Smeyers et al. 2021).

Of the three outliers found by two methods, one was located in an intron of the gene *MAF*, a transcription factor which has been found to play key roles during eye formation in birds, especially lens development (Reza and Yasuda 2004). The other two were in intergenic regions, where one was 25 kbp within the gene *KANK1* and the other within 8 kbp of a long non-coding RNA gene. *KANK1* belongs to the Kank protein family, and functions as a regulator of actin polymerization, actin stress fibre formation, and cell migration (Kakinuma et al. 2009). Kank genes are involved in vascular vessel development in vertebrates (Hensley et al. 2016). Long non-coding RNAs may regulate gene expression at pre-transcriptional, transcriptional and post-transcriptional levels, as well as contribute to epigenetic regulation (Chen et al. 2020; Robinson et al. 2020), and have been found for instance to play an important role in the avian immune system (Chen et al. 2020). However, we cannot determine the effects of the long non-coding RNA genes associated with our outliers, and we can only speculate about the significance of the above-mentioned genes to the reed warbler populations.

We looked closer at the allele frequencies of the four outliers found by at least two of the three genome scan methods (Table 9). We see that for the outlier close to *C9orf72*, Turkey has the most distinct allele frequency. For the outlier close to *KANK1*, Finland and Norway have relatively balanced allele frequencies, whereas one allele is fixed in the other populations. Finland is the most distinct for the outlier close to the long non-coding RNA gene, *LOC107053979*, and Germany is the most distinct for the outlier in *MAF* (Table 9).

**Table 9.** Allele frequencies of the first allele of the four outliers found by at least two of the three methods: BayeScan, Outflank and BayeScEnv for each population, sorted from highest to lowest latitude. Listed is the closest gene to the outlier.

| Country | C9orf72 | KANK1 | LOC107053979 | MAF |
| --- | --- | --- | --- | --- |
| Finland | 1 | 0.67 | 0.50 | 0.69 |
| Norway | 1 | 0.75 | 1 | 0.86 |
| Germany | 1 | 1 | 1 | 0.36 |
| Czech | 1 | 1 | 0.88 | 0.50 |
| Slovakia | 0.83 | 1 | 0.94 | 0.94 |
| France | 1 | 1 | 1 | 1 |
| Croatia | 1 | 1 | 0.83 | 0.71 |
| Italy | 0.95 | 1 | 0.86 | 0.75 |
| Turkey | 0.19 | 1 | 1 | 1 |
| Malta | 1 | 1 | 1 | 1 |

With BayeScEnv, we ran each bioclimatic predictor separately. Of the 38 outliers we found in total, 31 were found only by a single predictor (Supplementary table 8), and 7 were found by two or more predictors. One outlier was found by all six predictors, and one was found by all predictors except annual mean temperature (A.Temp), namely the two above-mentioned outliers associated with the genes *KANK1* and *C9orf72*, respectively (Supplementary table 7).

In a scatterplot of the first two principal component axes from a PCA of our 49 outliers, Malta is clearly separated on axis 1, with Norway and Finland at the opposite end (Figure 3C), which explains ∼13% of the variance in the dataset. There is a gradual south-to-north trend following axis 1 leftwards (Figure 3C). On axis 2, Turkey and an individual from Slovakia is on the opposite end from Norway and Malta.

We also searched for signs of recent selective sweeps throughout the genome by looking at haplotype homozygosity. Looking at -log_10_ p-value of iHS per chromosome for all populations combined, we see that the largest chromosomes (those >20 Mb, and especially those >50 Mb) have high peaks on either end of the chromosome (Figure 4). Most of the smaller chromosomes have many peaks spread across the whole chromosome. We set the threshold for being marked as an outlier quite high (-log_10_ p-value = 4, signifying a p-value of 0.0001), as a conservative measure since there were many high values. The pattern of outliers is for the most part consistent with the pattern of recombination rates in bird genomes, as recombination rate increases drastically towards chromosome ends in the macrochromosomes, and are uniformly high in microchromosomes (Backström et al. 2010). The high recombination rate may decrease the accuracy of statistical phasing (Weng et al. 2014), which in turn may possibly affect iHS results.

**Figure 4.**
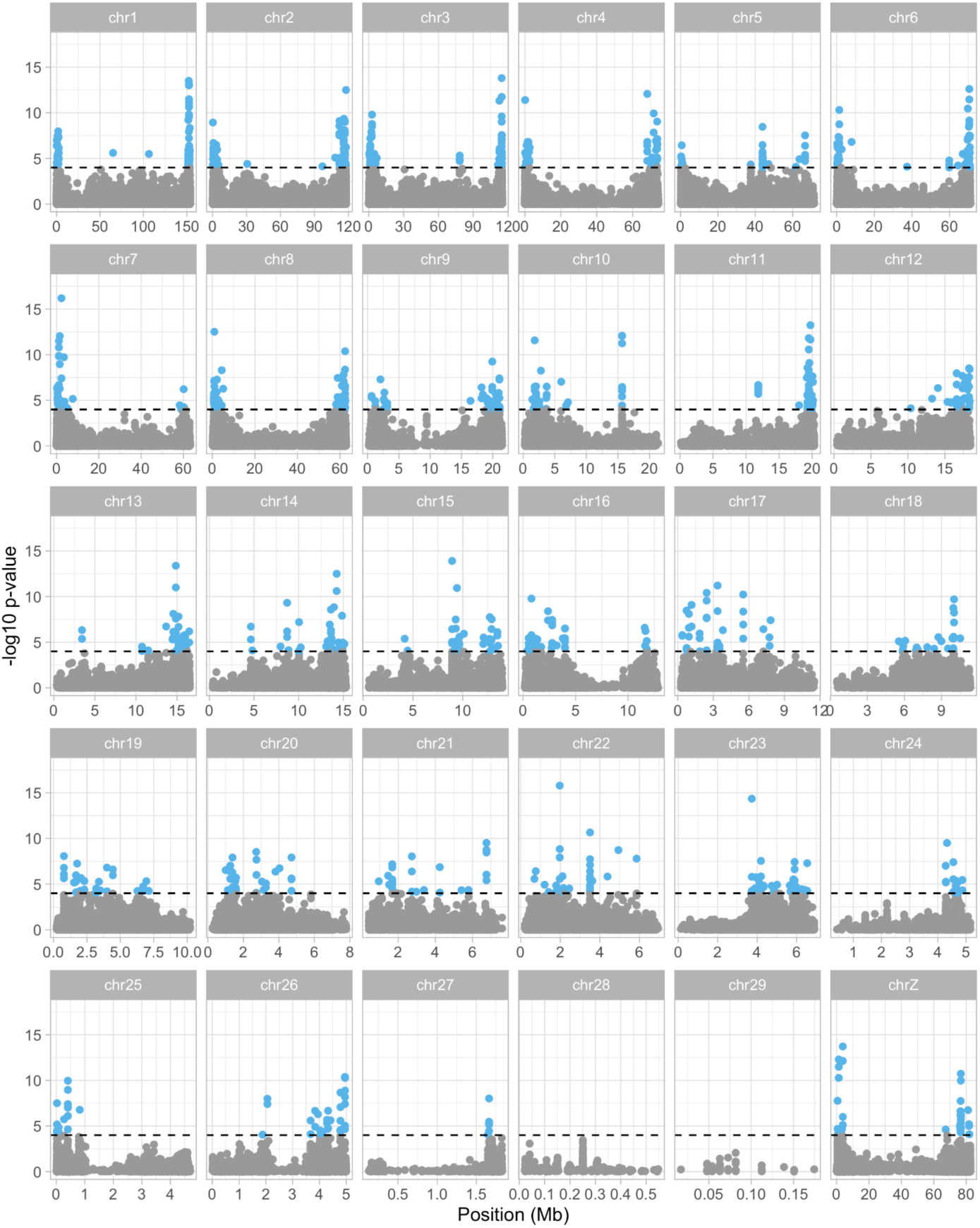
-Log10 p-values of iHS per chromosome for all populations combined. The stapled horizontal line is the threshold for being marked as an outlier (-log10 p-value = 4), and loci above this threshold are coloured light blue.

To investigate whether any of the outliers found by BayeScan, BayeScEnv and Outflank are associated with regions that may have undergone a selective sweep, we checked their iHS and average absolute XP-EHH values, along with the -log_10_ p-values. To see if the average iHS -log_10_ p-value of our genome scan outliers deviates significantly from a set of random loci, we sampled 43 random loci 10,000 times (not 49 since there were missing values for six outliers) and calculated the average iHS - log_10_ p-value for each set. On the histogram, the average value for the genome scan outliers is to the right of the 95th percentile, although the confidence interval is large (Figure 5; average = 0.67, 95% CI = 0.39–0.95). 15 of 43 outliers are above the 95th percentile for the random sets of loci. One of the outliers is a clear iHS outliers (-log_10_ p-value > 4, corresponding to p-value < 0.0001): the outlier on chromosome Z close to the gene *KANK1*, found by both BayeScan and BayeScEnv (all predictors for BayeScEnv). Another three outliers have iHS -log_10_ p-values of over 2 (p-values < 0.01); one located in an intron of the gene *HPCAL1*, and two in intergenic regions closest to long non-coding RNA genes. *HPCAL1* is a member of neuron-specific calcium-binding protein family found in the retina and brain, and may be involved in the regulation of rhodopsin phosphorylation and neuronal signalling in the central nervous system.

**Figure 5.**
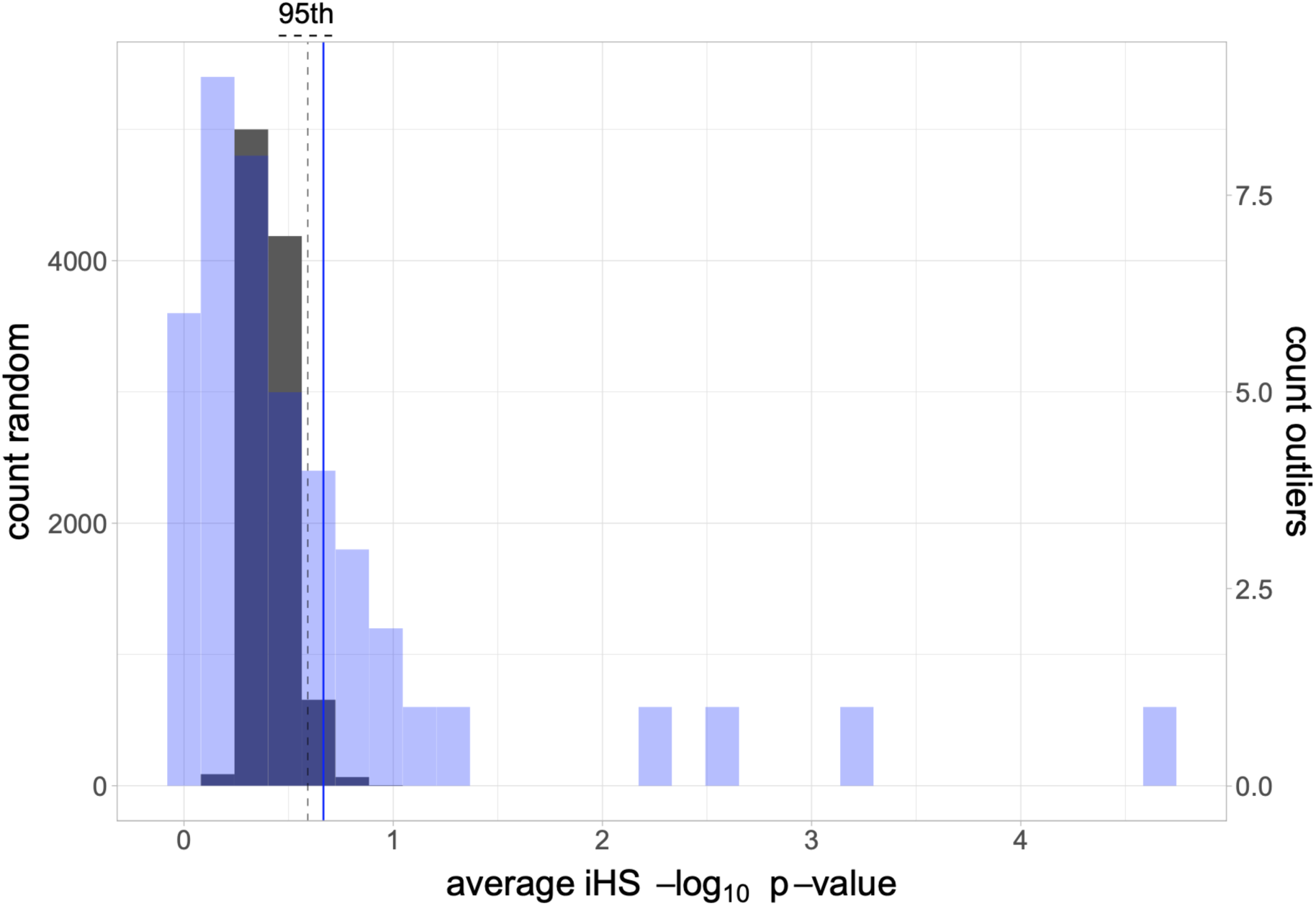
Histogram of average iHS -log10 p-values of 43 random loci 10,000 times (histogram in black colour and count on left y-axis), with the 95th percentile as a black stapled line. Overlaid is a histogram of iHS -log10 p-values of the genome scan outliers (N = 43, since there were six missing values) in blue colour, with count on the right y-axis, and the average values as a blue line.

We also checked if the XP-EHH -log_10_ p-values for our outlier loci deviate significantly from random sets, so we sampled 48 random loci 10,000 times (one outlier was missing from XP-EHH analyses because it was on a smaller scaffold) and calculated the average XP-EHH -log_10_ p-value for each set. The average XP-EHH -log_10_ p-value of our outliers is 0.43, which is lower than the 95th percentile for the random sets (Supplementary figure 5). Only one of the 2160 pairwise comparisons for the outlier loci has a -log_10_ p-value over 4 (p-value < 0.0001), and 47 (2.18%) have -log_10_ p-values over 2 (p-value < 0.01), so there seems to be a low correlation between outlier status in genome scans and significance in XP-EHH. The highest XP-EHH -log_10_ p-value is between Croatia and Slovakia on an outlier on chromosome 11 located in the intron of the gene *ARID3B*. This outlier was found by BayeScEnv and associated with precipitation seasonality. *ARID3B* belongs to a family of AT-rich interaction domain (ARID) proteins that are involved in for instance embryonic patterning, cell cycle control and chromatin remodelling.

Finally, using Hi-C data, we wanted to explore the significance of interactions between the genome scan outliers and transcription start sites of genes, specifically to see if the fraction of significant interactions for the outlier loci deviates from the fraction of significant interactions for sets of random loci. The fraction of significant interactions for the outliers is lower than average for the random sets, but is not significantly different (Figure 6).

**Figure 6.**
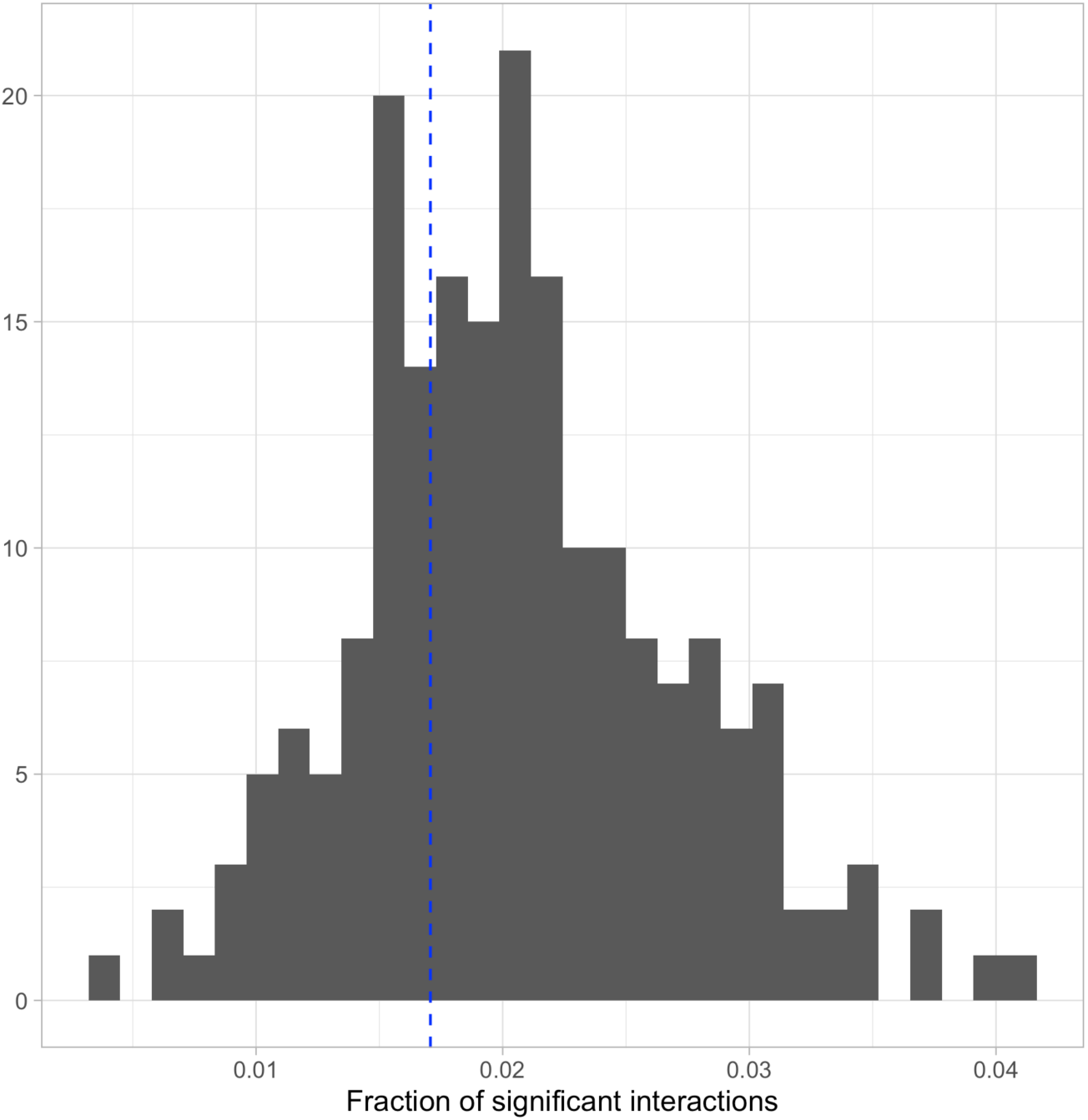
Histogram showing the fractions of significant interactions between SNPs and transcription start sites of genes, using Hi-C data, for 49 random loci sampled 200 times. The fraction of significant interactions for the 49 outlier loci from genome scans is marked as a blue stapled line.

## Discussion

Like many other contemporary species, the reed warbler is currently expanding its range northwards as a response to climate change, but the reed warbler has also recently expanded its range southwards, following habitat restoration near the southern range edge. These two range edges vary drastically climatically, and the two range expansions, different in direction and scale, provide a unique opportunity to study the genomic consequences of range expansion. In this study, we used RAD-seq to investigate population divergence and genetic patterns in European populations of the reed warbler following their recent northern range expansion and southern colonisation, and to look for signs of selection or environmental adaptation.

### Low genetic differentiation, and no apparent loss in genetic diversity after range expansion

We do not detect any substantial loss in genome-wide diversity in either the northern range expansion front (Finland and Norway), or in the newly established population at the southern range edge (Malta). Furthermore, we find that genetic structure is weak among the populations studied. Range expansion should in theory cause decreased genetic diversity at the range edge and increased population structure due to repeated founder events and strong genetic drift at the expansion front, unless gene flow remains high (Ray et al. 2003; Excoffier 2004; Excoffier et al. 2009). In the reed warbler, site fidelity is relatively high (Paradis et al. 1998; Ceresa et al. 2016), and individual reed warblers have been shown to build nests within one metre of the nest they built the year before (Long 1975). However, reed warblers are also capable of long-distance dispersal events (Paradis et al. 1998; Ceresa et al. 2016), and especially juveniles may disperse far away from their natal site (Bulyuk et al 2000; Mukhin 2004). Long-distance dispersal is thought to be an important factor in preserving genetic diversity at range edges and minimising genetic structuring during range expansion (Ray and Excoffier 2010, Berthouly-Salazar et al. 2013, Engler et al. 2016).

On a finer scale, we find that Malta is the most genetically distinct from the other populations, though pairwise global F_ST_ values are still low (< 0.04), and the populations at the northern range edge (Finland and Norway) separates from the remaining core populations. Of the populations studied, Malta is the youngest and the smallest. In the mid-1990’s, Is-Simar nature reserve on Malta had been fully restored, and reed warblers had just begun breeding there. Today the nature reserve contains the highest number of breeding pairs in Malta, but the area is still only about 5 hectares in total, and supports about 5-8 pairs on a consistent basis. We find that several pairs of the Maltese individuals have high relatedness scores, indicating close kinship, which may not be surprising considering the limited population size, as well as a high recruitment rate (Sætre et al. 2017). Perhaps more surprising is that there was no significant difference between Malta and the other populations in average nucleotide diversity, expected and observed heterozygosity or F_IS_ at the genome-wide level. This contradicts a previous finding that the Maltese population had a significantly higher F_IS_ than other European populations (Sætre et al. 2017). This may be due to differences in sampling design, since the previous study used microsatellites instead of RAD-seq. Microsatellites have a much higher mutation rate and typically produce higher heterozygosity estimates than RAD loci (Sunde et al. 2020). Depending on the ratio of observed and expected number of heterozygotes, F_IS_ estimates may be lower for microsatellites than for RADs, but may also be higher (Hodel et al. 2017). Missing data and allelic dropout in RAD-seq data have been found to be positively correlated with F_IS_ estimates (Marandel et al. 2020), and it is possible that RAD-seq or the subsequent filtering of SNPs may overestimate heterozygosity, if markers with relatively high frequency of both alleles are kept. Alternatively, it is possible that the high mutation rate in microsatellites allows for earlier detection of possible inbreeding occurring in Malta.

Although none of the populations differ significantly from the others, Norway have the lowest number of observed heterozygotes, and amongst the highest F_IS_ values. The heterozygosity values for Finland could be slightly inflated by a small batch effect, but accounting for that, they are most likely not lower than the values for Norway. Reed warblers colonised Finland in the 1920s (Järvinen and Ulfstrand 1980), but were first recorded breeding in Norway in 1947 (Røed 1994), making Norway the second youngest of the populations studied. The number of breeding pairs is likely still lower in Norway than in Finland, as it was around 20 years ago (Stolt 1999), but since the numbers continue to increase, the danger of inbreeding should be low in both of the northern range populations. Although the range expansion has been at least partially independent in Norway and Finland, there is very little genetic differentiation between these two populations, and they cluster away from the other populations.

### Isolation by environment stronger than isolation by distance

During rapid range expansion, patterns of isolation by distance may be expected if gene flow decreases with geographic distance. In this study, we find that there is significant isolation by distance from a simple Mantel test. However, in a multivariate regression analysis, looking at both geographic distance and environmental disparity together, geographic distance have a low unique contribution in explaining variance in genetic differentiation between populations. In species with low dispersal, isolation by distance is likely to be a consequence of range expansion, especially if the habitat is fragmented (Assis et al. 2013), but a lack of strong isolation by distance has been found in species capable of long-distance dispersal (Berthouly-Salazar et al. 2013; Heppenheimer et al. 2018). A previous study of the reed warbler found significant, but uneven levels of isolation by distance throughout the range (Procházka et al. 2011). However, isolation by distance only accounted for a small part of the genetic differences among populations (Procházka et al. 2011), and no climatic variables were investigated at that point.

In this study, the genetic differentiation between the reed warbler populations is due mainly to differences in climatic variables, rather than geographic distance. Precipitation seasonality have the single largest effect on genetic differentiation in our populations, explaining by itself 34% of the variance. In comparison, geographic distance explain only 4% of the variance in genetic differentiation. Malta is the most genetically distinct population, and clearly has the highest precipitation seasonality, meaning the amount of precipitation throughout the year fluctuates the most of all populations. Compared to Finland and Norway, which are genetically most differentiated from Malta, Malta gets lower annual precipitation, but slightly more precipitation in the wettest, coldest quarter. In the warmest, driest quarter on the other hand, Malta gets significantly less, and in the driest month, there is practically no precipitation. Like precipitation seasonality, different temperature variables vary quite a lot between populations, and are likely to have an effect on gene flow. We find that the first principal component of all chosen temperature and precipitation variables explain by itself 23% of the variance in genetic differentiation.

Gene flow between similar environments may facilitate local adaptation during range expansion by introducing new alleles that are locally beneficial (Sexton et al. 2011). Isolation by environment has been found to better explain the patterns of genetic differentiation than isolation by distance, and to have positive effects on fitness at range edges (Kottler et al. 2021). Immigrants from climatically different environments may be selected against or mating could be non-random (Sexton et al. 2014). We would have liked to get a direct estimation of migration rates or gene flow among populations, by for instance using BayesAss (Wilson and Rannala 2003), but this method is not reliable when divergence among populations is very recent or population structure is low (F_ST_ < 0.10) (Meirmans, 2014). The low population structure between reed warbler populations suggests that the populations had not had time to diverge yet, or that gene flow is high, but it seems gene flow is partially impeded by adaptation to local climatic conditions.

The range expansion analysis confirms the reed warbler’s recent range expansion northwards into Fennoscandia (Järvinen and Ulfstrand 1980; Røed 1994; Stolt 1999; Brommer et al. 2012), and the strongest signals identified Norway and Finland as furthest away from the estimated point of origin. The reed warbler had a post-glacial expansion after surviving the Last Glacial Maximum in several different refugia (Arbabi et al. 2014), so we cannot know the exact point of origin for the reed warbler populations studied here. Although Malta is the most recently established population, the mechanism of the establishment is different from the northwards range expansion. It is likely that the rapid colonisation of Malta consisted mainly of reed warblers originating from southern populations, i.e., a ghost source population not included in our sampling, which would explain why it is not picked up by the range expansion analysis.

### Signatures of selection and the role of climate as a driver of local adaptation

We find that there is low correlation between genetic differentiation between range edges and differentiation within the range core, suggesting novel genetic variation has arisen at the range edges to some extent. Several loci are outliers in terms of being highly differentiated between populations, most of which also covary with climatic differentiation. Of the six different (not highly correlated) temperature and precipitation variables tested, all of them are associated with at least one outlier. A higher number of outliers are associated with precipitation variables than temperature variables, although the difference is not very large (19 outliers are associated with temperature variables, 33 outliers are associated with precipitation variables). A PCA of the outlier loci separates the northern and southern range edge on the first axis, reflecting the population structure in the full dataset. It is likely that both temperature and precipitation are important drivers of local adaptation in the reed warbler, perhaps especially at the range edges.

The reed warbler is a wetland bird breeding mainly in *Phragmites* reed beds, and both temperature and precipitation may affect reed growth, structure and biomass (Engloner 2009), as well as the availability of insects, as many develop in aquatic or moist habitats (Frampton et al. 2000; Jiménez et al. 2018). These factors may in turn affect things like timing of nest building, egg laying, incubation period and clutch size. Newly emerged reeds provide better protection from predation, so when conditions are favourable for reed growth, reed warblers may have fewer nest losses. This may be part of the explanation for why reed warblers have begun breeding earlier following warmer spring temperatures (Halupka et al. 2008). Prolonged periods of drought have been shown to negatively affect productivity in reed warblers in Spain, likely due to decreased availability of insects (Jiménez et al. 2018). However, egg laying and nestling mortality may be negatively affected by heavy rain in the breeding season (Halupka et al. 2008; Vafidis et al. 2016). Both precipitation and temperature have been shown to drive selection in reed warblers, for example in wing length (Nowakowski 2000), migration strategy (Chamorro et al. 2019) and egg coloration (Avilés et al. 2007).

One of the genome scan outliers correlate with all precipitation and temperature variables tested in BayeScEnv, and is also marked as an outlier by BayeScan. In all populations except Finland and Norway the birds are fixed for one allele. In these northern populations, however, a second allele is also segregating at an intermediate frequency. This private allele present in Finland and Norway may be a new mutation, which has arisen either independently in the two populations, or in one and transferred via gene flow. Another possibility is that this variant is present in the range core, but at a very low frequency which has not been captured by our sampling, and that the variant has been picked up by selection during founder events. According to the iHS analysis, the outlier is involved in a recent selective sweep. However, this should be interpreted with caution, since it is located near the start of the chromosome, and all of the largest chromosomes have high iHS peaks at the chromosome ends. This pattern may be due to the high recombination rates at chromosome ends in the macrochromosomes in birds (Backström et al. 2010), which may cause statistical phasing to be less accurate (Weng et al. 2014). However, selection may also be more effective in high-recombination regions, and adaptive substitutions have been found to appear more frequently in these areas (Gossmann et al. 2014).

Although the outlier is located in an intergenic region, it is close to the gene *KANK1*, which belongs to the Kank protein family, and functions as a regulator of actin polymerization, actin stress fibre formation, and cell migration (Kakinuma et al. 2009). Kank genes are involved in vascular vessel development in vertebrates (Hensley et al. 2016), and may play a role in the well-vascularized uterine tissue required for viviparity (Recknagel et al. 2021), but it is unclear what the exact function of this gene is in birds. Furthermore, we cannot be sure if this gene is the actual target of selection in the reed warbler.

All of the other outliers are located either in introns, or in intergenic regions. Since none of our outliers were found within an exon, we cannot be sure whether they directly impact gene expression, nor what significance they have for our reed warbler populations. We find that our outliers on average have significantly higher iHS -log_10_ p-values than random sets of loci, meaning some of them seem to be part of recent selective sweeps. The high average value is probably because of the few strong signals, so many outlier loci may be under selection and yet not in the process of undergoing a sweep. Perhaps the recent character of the range expansion makes them difficult to detect, and possibly this pattern may change in the future. Overall, we find many regions throughout the genome putatively involved in recent selective sweeps with the iHS analysis. However, more research is needed to examine the effect of recombination rate on the efficacy of statistical phasing, and whether this affects the iHS results.

Using Hi-C data, we calculated the significance of interactions between the genome scan outliers and transcription start sites of genes and compared this to sets of random loci. As far as we know, this has not been done before in this context, but we wanted to explore the hypothesis that genome scan outliers have a higher than random fraction of significant interactions with transcription start sites, as this could facilitate linked selection and selective sweeps. There was no significant difference between our outlier loci and the random loci, so it does not seem to be the case here.

## Conclusion

In this study, we investigated the genomic consequences of range expansion in the reed warbler. This represents a unique case, where the range expansion has occurred in two directions following different human-induced expansion processes: one directly induced by humans (habitat restoration) and one indirectly (climate change). We find no apparent loss in genetic diversity at either range edge, and low genetic differentiation among the reed warbler populations. However, gene flow is partially impeded by adaptation to local climatic conditions. Temperature variables, and especially precipitation variables induce significant isolation by environment, and correlate with several loci putatively under selection. Local adaptation to climatic conditions may have facilitated the reed warbler’s range expansion both northwards and southwards, and may thus be crucial for their success.

## Acknowledgements

We thank Ingvild Myhre Ekeberg for assisting with field work. We also thank Glenn-Peter Sætre for assembling the panel figures. Furthermore, we are grateful to the Natural History Museum in Oslo for supplying additional samples. Funding was granted by the Norwegian Research Council to Fabrice Eroukhmanoff and a PhD grant to Camilla Lo Cascio Sætre from the University of Oslo.

## Notes

### Competing Interest Statement

The authors have declared no competing interest.

