## Supplementary methods, Supplementary Tables 1-7, Supplementary Figures 1-5 for "Genomic consequences of range expansion and colonisation in the reed warbler (*Acrocephalus scirpaceus*)"

### Supplementary materials

#### Supplementary methods

We extracted the 19 bioclimatic predictors (30 s) from WorldClim (Hijmans et al. 2005) and used the raster package v3.4-13 (Hijmans 2021) in R to upload them as RasterLayers and extract points from the coordinates of each site (Supplementary table 1). The program was not able to extract values for the bioclimatic predictors for the coordinates for Storminnet in Finland, so for the two individuals from this site, we used the values for the closest site, Kilpilahti. These sites are very close (Storminnet: 60.32, 25.59; Kilpilahti: 60.32, 25.54).

**Supplementary table 1.** Overview of all sites sampled in this study, including coordinates, sample size (N) and six selected environmental variables from WorldClim. Longitude and latitude are presented as decimal degrees (rounded to the nearest hundredths). A.Temp = annual mean temperature (°C). Temp.S = temperature seasonality (standard deviation * 100, °C). M.Temp = maximum temperature of the warmest month (°C). A.Prec = annual precipitation (millimetres). Prec.S = precipitation seasonality (CV, percentage). C.Prec = precipitation of the coldest quarter (millimetres).

| **Country** | **Site** | **Long.** | **Lat.** | **N** | **A.Temp** | **Temp.S** | **M.Temp** | **A.Prec** | **Prec.S** | **C.Prec** |
|---|---|---|---|---|---|---|---|---|---|---|
| Croatia | Neretva | 43.06 | 17.61 | 9 | 16.52 | 722 | 32.00 | 1001 | 31.96 | 302 |
| Czech | Lužice | 48.86 | 17.08 | 5 | 9.61 | 765 | 25.20 | 629 | 41.56 | 96 |
| Czech | Osík | 49.85 | 16.28 | 5 | 7.91 | 721 | 23.40 | 527 | 48.18 | 72 |
| Finland | Bjurböleviken | 60.34 | 25.72 | 2 | 4.56 | 847 | 21.80 | 666 | 28.07 | 150 |
| Finland | Kilpilahti | 60.32 | 25.54 | 4 | 4.72 | 847 | 22.00 | 674 | 28.59 | 154 |
| Finland | Kodderviken | 60.34 | 25.60 | 4 | 4.53 | 848 | 21.80 | 671 | 28.48 | 152 |
| Finland | Källsundet | 60.34 | 25.70 | 2 | 4.57 | 847 | 21.80 | 666 | 27.79 | 151 |
| Finland | Storminnet | 60.32 | 25.59 | 2 | 4.72 | 847 | 22.00 | 674 | 28.59 | 154 |
| France | Trunvel | 47.90 | -4.36 | 9 | 12.14 | 361 | 20.60 | 1011 | 36.37 | 311 |
| Germany | Diergarten | 51.23 | 6.09 | 8 | 9.89 | 564 | 22.50 | 738 | 13.31 | 180 |
| Germany | Mohrhof | 49.67 | 10.85 | 2 | 8.61 | 682 | 23.70 | 707 | 16.92 | 166 |
| Italy | Lago Salso | 41.54 | 15.89 | 13 | 16.17 | 632 | 30.80 | 442 | 24.43 | 116 |
| Malta | Is-Simar | 35.95 | 14.38 | 11 | 18.90 | 510 | 30.60 | 516 | 83.23 | 177 |
| Norway | Hellesjøvannet | 59.74 | 11.46 | 4 | 5.20 | 754 | 20.80 | 786 | 27.46 | 152 |
| Norway | Hærsetersjø | 59.65 | 11.38 | 3 | 5.40 | 740 | 20.70 | 796 | 27.36 | 157 |
| Norway | Kragtorpvika | 59.67 | 11.48 | 8 | 5.33 | 744 | 20.60 | 787 | 27.16 | 155 |
| Slovakia | Trnava | 48.36 | 17.55 | 10 | 9.83 | 779 | 26.40 | 625 | 31.16 | 111 |

| | | | | | | | | | | |
|---|---|---|---|---|---|---|---|---|---|---|
| Turkey | Mogan | 39.75 | 32.79 | 8 | 10.93 | 832 | 29.50 | 383 | 41.46 | 123 |

**Supplementary table 2.** SNP count after different filtering steps.

| Filtering step | Filter applied | SNP count |
|---|---|---|
| 1 | Locus present in ≥ 80% of individuals in a population | 594,277 |
| 2 | 10 ≤ Coverage depth ≤ 80 | 522,533 |
| 3 | Genotype present in ≥ 80% of samples | 90,558 |
| 4 | Minor allele frequency (MAF) > 0.05 | 22,862 |
| 5 | Keep one SNP per RAD locus | 10,167 |
| 6 | Cut SNPs in linkage disequilibrium | 9,320 |
| 7 | Cut SNPs in Hardy-Weinberg disequilibrium | 9,236 |

**Supplementary table 3**. Differences between the five control samples run in both 2017 and in 2021, in mean depth (Depth), frequency of missing data (Miss.) and heterozygosity (Het.). The samples are single individuals from Czech (CZ), France (FR), Italy (IT), Norway (NO) and Turkey (TK).

| Sample | Depth 2017 | Depth 2021 | Miss. 2017 | Miss. 2021 | Het. 2017 | Het. 2021 |
|---|---|---|---|---|---|---|
| CZ | 99.23 | 42.95 | 0.12 | 0.13 | 0.08 | 0.08 |
| FR | 44.28 | 42.46 | 0.17 | 0.13 | 0.06 | 0.08 |
| IT | 64.00 | 42.66 | 0.14 | 0.12 | 0.07 | 0.09 |
| NO | 36.16 | 18.45 | 0.19 | 0.14 | 0.05 | 0.08 |
| TK | 44.92 | 37.11 | 0.16 | 0.13 | 0.07 | 0.09 |

**Supplementary table 4**. Basic statistics within populations. Mean nucleotide diversity ($\pi$), mean observed heterozygosity ($H_O$), mean expected heterozygosity ($H_E$) and $F_{IS}$ is from the sumstats_summary file from Stacks *populations* using -r 0.80. Allelic richness (AR) is estimated with hierfstat using the VCF from the same *populations* run.

| Country | $\pi$ | AR | $H_O$ | $H_E$ | $F_{IS}$ |
|---|---|---|---|---|---|
| Finland | 0.100 | 1.352 | 0.088 | 0.096 | 0.039 |

| | | | | | |
|---|---|---|---|---|---|
| Norway | 0.095 | 1.334 | 0.060 | 0.092 | 0.122 |
| Germany | 0.098 | 1.336 | 0.066 | 0.092 | 0.094 |
| Czech | 0.102 | 1.357 | 0.078 | 0.097 | 0.072 |
| Slovakia | 0.108 | 1.399 | 0.069 | 0.102 | 0.129 |
| France | 0.093 | 1.323 | 0.063 | 0.087 | 0.087 |
| Croatia | 0.098 | 1.347 | 0.075 | 0.092 | 0.067 |
| Italy | 0.098 | 1.347 | 0.071 | 0.094 | 0.091 |
| Turkey | 0.101 | 1.362 | 0.076 | 0.094 | 0.071 |
| Malta | 0.095 | 1.323 | 0.066 | 0.091 | 0.090 |

**Supplementary table 5**. Range expansion statistics. Average pairwise values of the directionality index (Ψ) along with the average latitude of the population sampled, sorted from highest to lowest latitude.

| Population | Latitude | Average pairwise Ψ |
|---|---|---|
| Finland | 60.3 | 0.0293 |
| Norway | 59.7 | 0.0148 |
| Germany | 50.9 | -0.0030 |
| Czech | 49.4 | 0.0010 |
| Slovakia | 48.4 | -0.0012 |
| France | 47.9 | 0.0034 |
| Croatia | 43.1 | -0.0080 |
| Italy | 41.5 | 0.0053 |
| Turkey | 39.8 | -0.0250 |
| Malta | 35.9 | -0.0168 |

**Supplementary table 6**. Four outliers found by at least two of the three following methods: BayeScan (Ba), Outflank (Ou) and BayeScEnv (Be). The genomic positions in the reed warbler genome of the loci are noted, along with the number of base pairs (bp) to the closest gene and its corresponding gene name. The gene biotype is also noted, where prot means protein-coding and lncRNA means long non-coding RNA. BayeScEnv was run separately for six selected bioclimatic predictors: A.Temp = annual mean temperature, Temp.S = temperature seasonality, M.Temp = maximum temperature of the warmest month, A.Prec = annual precipitation, Prec.S = precipitation seasonality, and C.Prec = precipitation of the coldest quarter. The final column shows which of the BayeScEnv runs marked the locus as an outlier.

| Chr | Locus | Position | Closest gene bp | Gene Name | Biotype | Found by | BayeScEnv predictors |
|---|---|---|---|---|---|---|---|
| 4 | 65612233 | Intergenic | 8,020 | *LOC107053979* | lncRNA | Ba, Be | Temp.S |
| 10 | 5923709 | Intron | 0 | *MAF* | prot | Ba, Be | Temp.S |
| Z | 3368695 | Intergenic | 25,064 | *KANK1* | prot | Ba, Be | All |
| Z | 46111776 | Intergenic | 14,940 | *C9orf72* | prot | Ba, Ou, Be | All (-A.Temp) |

**Supplementary table 7**. Outliers found by BayeScEnv analysis run separately for each bioclimatic predictor. A.Temp = annual mean temperature. Temp.S = temperature seasonality. M.Temp = maximum temperature of the warmest month. A.Prec = annual precipitation. Prec.S = precipitation seasonality. C.Prec = precipitation of the coldest quarter. Each row shows the total number of outliers found by a predictor, and of these, how many were found only by that predictor. A total of 31 outliers were found by a single predictor, and another 7 were found by two or more predictors, totalling 38 unique outliers.

| Bioclimatic variable | Total N outliers | Unique N outliers |
|---|---|---|
| A.Temp | 1 | 0 |
| Temp.S | 11 | 8 |
| M.Temp | 7 | 3 |
| A.Prec | 8 | 2 |
| Prec.S | 9 | 6 |
| C.Prec | 16 | 12 |

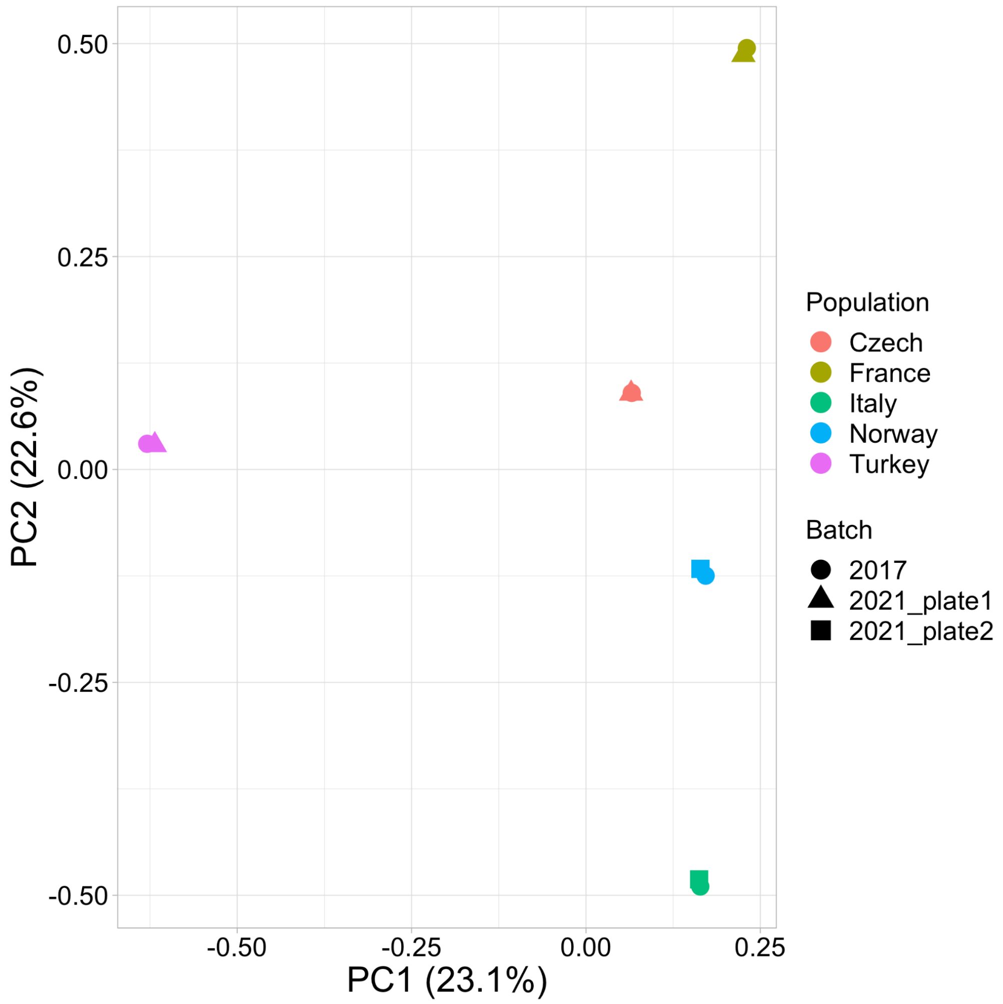


**Supplementary figure 1**. Controlling for batch effects. The first two principal axes of a PCA analysis with the five control samples which had been run both in 2017 on a single plate with all populations except Finland, and in 2021 split over two plates, with the Finnish samples. The control samples were added to the 2021 plates along with the Finnish samples, to check for possible batch effects. The colour corresponds to the population of the sample, and the shape corresponds to the plate the sample was run in. In parentheses, the percentage of variance explained by each axis.

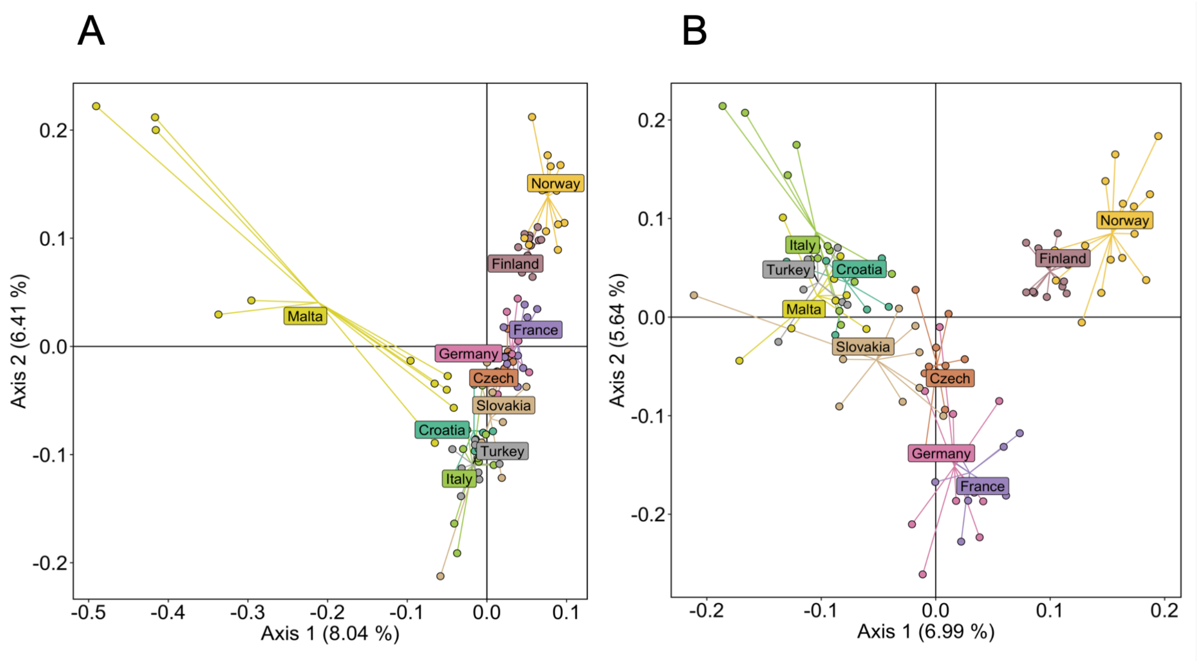


**Supplementary figure 2**. Exploring populations structure with PCA. A: The first two principal axes of a PCA analysis with all individuals. In parentheses, the percentage of variance explained by each axis. B: The first two principal axes of a PCA analysis with three individuals from Malta removed. These individuals were part of a trio and duo that clustered far away from the other samples, and had high relatedness ($A_{jk} > 0.3$) amongst each other. In parentheses, the percentage of variance explained by each axis.

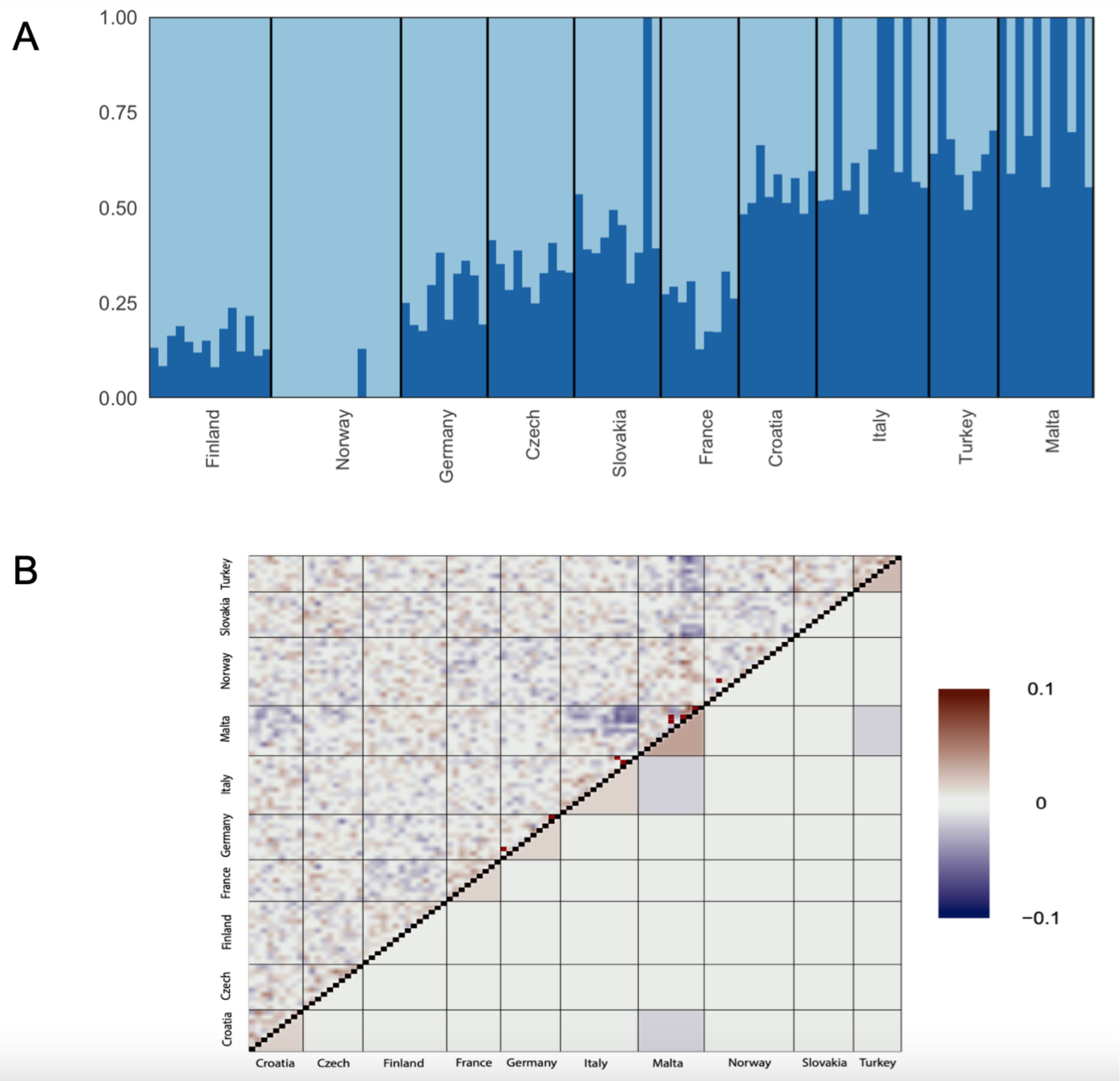


**Supplementary figure 3**. Assessing the fit of an admixture model with assumed number of populations (K) = 2. A: Admixture plot of K = 2, where the ancestral fractions are represented by dark blue and light blue colour. The 10 populations are sorted from highest to lowest latitude going right. B: Plot of correlation of residuals for K = 2 from evalAdmix. The upper diagonal shows the correlation of residuals between individuals and the lower diagonal shows mean residual correlation within and between populations. Correlation values above and below zero on the colour scale are set to dark red and dark blue, respectively.

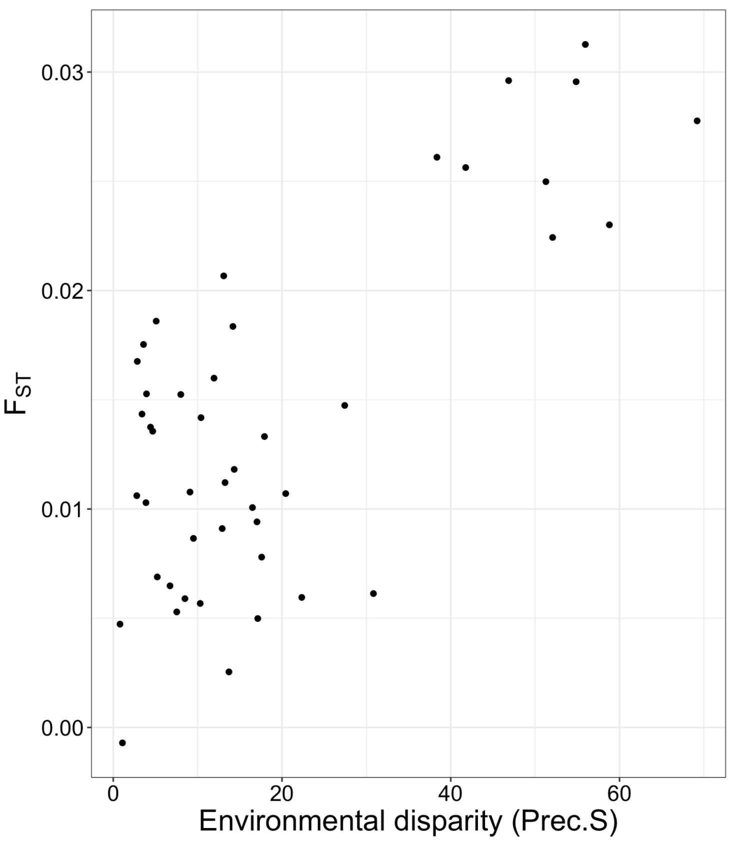


**Supplementary figure 4**. Isolation by environment. Pairwise $F_{ST}$ against environmental disparity in precipitation seasonality (Prec.S) between each population pair.

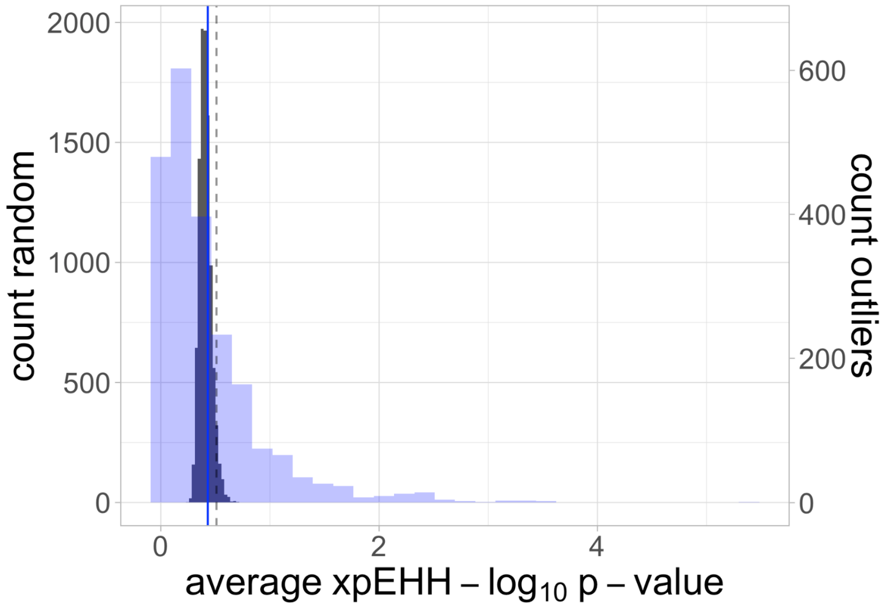


**Supplementary figure 5**. Histogram of average XP-EHH -$log_{10}$ p-values of 48 random loci 10,000 times (histogram in black colour and count on left y-axis), with the 95th percentile as a black stapled line. Overlaid is a histogram of XP-EHH -$log_{10}$ p-values of the genome scan outliers (N = 48, since there was one missing value) in blue colour, with count on the right y-axis, and the average values as a blue line.
